# Calcium-permeable AMPA receptors mediate timing-dependent LTP elicited by low repeat coincident pre- and postsynaptic activity at Schaffer collateral-CA1 synapses

**DOI:** 10.1101/719633

**Authors:** Efrain A. Cepeda-Prado, Babak Khodaie, Gloria D. Quiceno, Swantje Beythien, Elke Edelmann, Volkmar Lessmann

## Abstract

High-frequency stimulation induced long-term potentiation (LTP), or low frequency stimulation induced LTD are considered as cellular models of memory formation. Interestingly, spike timing-dependent plasticity (STDP) can induce equally robust timing-dependent LTP (t-LTP) and t-LTD in response to low frequency repeats of coincident action potential (AP) firing in presynaptic and postsynaptic cells. Commonly, STDP paradigms relying on 25-100 repeats of coincident AP firing are used to elicit t-LTP or t-LTD, but the minimum number of repeats required for successful STDP is barely explored. However, systematic investigation of physiologically relevant low repeat STDP paradigms is of utmost importance to explain learning mechanisms *in vivo*. Here, we examined low repeat STDP at Schaffer collateral-CA1 synapses by pairing one presynaptic AP with either one postsynaptic AP (1:1 t-LTP), or a burst of 4 APs (1:4 t-LTP) and found 3-6 repeats to be sufficient to elicit t-LTP. 6x 1:1 t-LTP required postsynaptic Ca^2+^ influx via NMDARs and L-type VGCCs and was mediated by increased presynaptic glutamate release. In contrast, 1:4 t-LTP depended on postsynaptic metabotropic GluRs and ryanodine receptor signaling, and was mediated by postsynaptic insertion of AMPA receptors. Unexpectedly, both 6x t-LTP variants were strictly dependent on activation of postsynaptic Ca^2+^-permeable AMPARs but were differentially regulated by dopamine receptor signaling. Our data show that synaptic changes induced by only 3-6 repeats of mild STDP stimulation occurring in ≤ 10 s can take place on time scales observed also during single trial learning.

## Introduction

Long-term potentiation (LTP) and long-term depression (LTD) of synaptic transmission can be observed in response to repetitive activation of synapses and are believed to represent cellular models of learning and memory processes in the brain (see e.g., Bi and Poo, 1998; Bliss and Cooke, 2011; Malenka and Bear, 2004). While LTP leads to a stable enhancement of synaptic transmission between connected neurons, LTD yields a long-lasting decrease in synaptic responses. Depending on the time frame that is investigated, LTP as well as LTD can be divided into an early phase lasting roughly 1h and a late phase that starts 2-3h after induction of the synaptic change. While early LTP is mediated by posttranslational modifications, late LTP was found to depend on synthesis of new proteins (Lynch, 2004; but see Wang et al., 2016). LTP was initially discovered using long-lasting high frequency stimulation of glutamatergic synapses in the mammalian hippocampus (Bliss and Lomo, 1973), a brain region essential for encoding episodic memory (see e.g., Eichenbaum, 1999; Fletcher et al., 1997; Squire and Zola, 1998; Tulving and Markowitsch, 1998) reviewed in (Tonegawa et al., 2018). While in these pioneering studies, LTP was recorded *in vivo* using extracellular field potential recordings (Bliss and Lomo, 1973), LTP is also observed in acutely isolated brain slices *ex vivo* and can be recorded in individual neurons using whole cell patch clamp recording techniques (reviewed e.g. in Herring and Nicoll, 2016; Lalanne et al., 2018; Pinar et al., 2017). Notably, LTP studies at the single cell level are essential to understand the biochemical and cellular mechanisms of LTP and LTD processes of a specific neuronal connection with defined postsynaptic target. To relate results from LTP measurements in acute slices *ex vivo* with learning processes it is important to use LTP induction protocols that resemble synaptic activation patterns also occurring during memory formation *in vivo* (compare Bittner et al., 2015; 2017; Otto et al., 1991), rather than paradigms involving tetanic synaptic stimulation or long-lasting artificial depolarization of postsynaptic neurons.

In this respect, spike timing-dependent plasticity (STDP) seems to represent an especially relevant protocol for induction of LTP (e.g., Bi and Poo, 2001; Caporale and Dan, 2008; Costa et al., 2017; Edelmann et al., 2017; Edelmann et al., 2014; Feldman, 2012; Markram et al., 2011). Here, bidirectional plasticity can be induced by repeated coincident activation of pre- and postsynaptic neurons, with forward pairing (i.e. presynaptic spike occurs several ms before the postsynaptic action potential; positive spike timing (+Δt)) yielding timing-dependent (t-) LTP, while backward pairing (postsynaptic spike occurs before presynaptic activation; -Δt) yields t-LTD. These protocols also fulfill the prerequisites for Hebbian synaptic plasticity (Caporale and Dan, 2008) that are widely accepted as fundamental requirements for synaptic plasticity. Compared to pairing protocols that induce LTP by combining a presynaptic tetanus with a postsynaptic depolarization (e.g., Meis et al., 2012), STDP relies on a small number of pre- and postsynaptic action potentials that are repeated at low frequency (<5 Hz).

Like memory formation *in vivo*, t-LTP in acute *ex vivo* brain slices is strongly controlled by neuromodulatory inputs, which can regulate the efficacy of induction paradigms to elicit plasticity (Edelmann and Lessmann, 2011, 2013; reviewed in Edelmann and Lessmann, 2018; Liu et al., 2017; Pawlak and Kerr, 2008; e.g., Seol et al., 2007). Such kind of neuromodulation can also bridge the temporal gap between synaptic plasticity and behavioral time scales for learning processes (Gerstner et al., 2018; Shindou et al., 2019), to connect synaptic effects to behavioral readouts. T-LTP was also reported to depend on brain-derived neurotrophic factor signaling (e.g., Edelmann et al., 2015; Lu et al., 2014; Mu and Poo, 2006; Pattwell et al., 2012; Sivakumaran et al., 2009). Still, these mechanistic studies on t-LTP employed STDP protocols depending typically on 25-100 repeats at ≤ 1 Hz that are unlikely to occur at synapses during memory formation *in vivo*. Therefore, in the present study we started out to determine the minimum number of repeats of coincident activation of pre- and postsynaptic neurons required for successful t-LTP induction. To this aim, we used low repeat variants of our recently described canonical (1:1 pairing: Edelmann and Lessmann, 2011) and burst STDP protocols (1:4 pairing: Edelmann et al., 2015; Solinas et al., 2019). Although, STDP protocols involving low repeat synaptic activation have been used previously to induce t-LTP (somatosensory cortex: Cui et al., 2015; Cui et al., 2016; visual cortex: Froemke et al., 2006; cultured hippocampal cells: Zhang et al., 2009), the underlying cellular mechanisms for its induction and expression remained elusive.

Our present study demonstrates that Schaffer collateral (SC)-CA1 t-LTP can be induced robustly by only three to six repeats of coincident pre- and postsynaptic spiking at 0.5 Hz. Moreover, our study reveals that, depending on specific STDP paradigms (i.e. 1:1 vs. 1:4) the low repeat protocols recruit distinct sources for postsynaptic Ca^2+^ elevation during induction of t-LTP, are mediated by distinct pre- and postsynaptic expression mechanisms, and are differentially regulated by dopamine receptor signaling. Together these data suggest that hippocampal SC-CA1 synapses can recruit multiple types of synaptic plasticity in response to low repeat STDP protocols for processing of information and memory storage in the hippocampus. Altogether, these distinct cellular STDP pathways might form the basis for the pluripotency of hippocampal functions in learning and memory.

## Material and Methods

### Preparation of hippocampal slices

Horizontal hippocampal slices (350 μm thickness) were prepared from 4 weeks old male wild type C57BL/6J (Charles River), BDNF^+/-^ or littermate control mice (Korte et al., 1995; all animals bred on a C57BL/6J background), according to the ethical guidelines for the use of animal in experiments, and were carried out in accordance with the European Committee Council Directive (2010/63/EU) and approved by the local animal care committee (Landesverwaltungsamt Sachsen-Anhalt).

Briefly, mice were decapitated under deep anesthesia with forene (Isofluran CP, cp-pharma, Germany) and the brain was rapidly dissected and transferred into ice-cold artificial cerebrospinal fluid (ACSF) cutting solution (125 mM NaCl, 2.5 mM KCl, 0.8 mM NaH_2_PO_4_, 25 mM NaHCO_3_, 25 mM Glucose, 6 mM MgCl_2_, 1 mM CaCl_2_; pH 7.4; 300-303 mOsmol/kg), saturated with 95% O_2_ and 5% CO_2_. Blocks from both hemispheres containing the hippocampus and the entorhinal cortex were sectioned with a vibratome (VT 1200 S, Leica, Germany). Slices were incubated for 35 min at 32°C in a handmade interface chamber containing carboxygenated ACSF cutting solution and then transferred to room temperature (∼21°C) for at least 1 hour before the recording started. Whole cell patch-clamp recordings were performed in submerged slices in a recording chamber with continuous perfusion (1-2 ml per min) of pre-warmed (30 ± 0.2°C) carboxygenated physiological ACSF solution (125 mM NaCl, 2.5 mM KCl, 0.8 mM NaH_2_PO_4_, 25 mM NaHCO_3_, 25 mM Glucose, 1 mM MgCl_2_, 2 mM CaCl_2_; pH 7.4; 300-303 mOsmol/kg). For all experiments, 100 µM Picrotoxin (GABA_A_ blocker) was added to ACSF solution. Epileptiform activity by activation of recurrent CA3 synapses was prevented by a cut between CA3 and CA1 subfields (compare Edelmann et al., 2015). To reduce the amount of inhibitors in some of the experiments (e.g. application of NASPM and IEM-1460) we used a micro-perfusion pump-driven solution recycling system (Bioptechs Delta T Micro-Perfusion pump high flow, ChromaPhor, Germany) to limit the volume of solution for incubation of slices (Meis et al., 2012). Both, NASPM and IEM were applied 15 min prior to STDP induction. The drugs were present during the whole experiment. Respective matched control experiments were performed under identical conditions to assure that the microperfusion recycling of ACSF alone did not affect t-LTP.

### Electrophysiological recordings

Whole cell patch-clamp recordings were performed on pyramidal neurons in the CA1 subregion of the intermediate hippocampus under visual control with infrared DIC-videomicroscopy (RT-SE series; Diagnostic instruments, Michigan, USA). The pipettes (resistance 5-7 MΩ) were filled with internal solution containing (in mM): 10 HEPES, 20 KCl, 115 potassium gluconate, 0-0.00075 CaCl_2_ added to the nominally Ca^2+^-free internal solution, 10 Na-phosphocreatine, 0.3 Na-GTP, and 4 Mg-ATP (pH 7.4, 285-290 mOsmol/kg). Cells were held at -70 mV in current clamp or voltage clamp (liquid junction potential of +10 mV for the combination of internal and external solutions was corrected manually) with an EPC-8 patch clamp amplifier (HEKA, Lamprecht, Germany). Extracellular stimulation of the Schaffer collateral (SC) fibers to generate an excitatory postsynaptic potential (EPSP, at 0.05Hz) was induced by glass stimulation electrodes (opening diameter: <15µm) filled with saline (resistance 0.7 – 0.9 MΩ) positioned in Stratum radiatum (SR) of the CA1 subregion. The stimulus intensity was adjusted to evoke responses with amplitudes of 4-7 mV corresponding to 30-50% of maximal EPSP amplitudes. Stimulus duration was set to 0.7 ms with intensities ranging between 90 to 700 µA.

### Induction of Spike timing-dependent plasticity

Spike timing-dependent plasticity (STDP) was induced by pairing of a single EPSP, generated by extracellular stimulation of SC, with a single action potential (AP) or with a burst of 4 APs (frequency 200 Hz) induced by somatic current injection (2 ms; 1 nA) through the recording electrode (Edelmann et al., 2015). Pairings of postsynaptic EPSP and APs were usually performed with a time interval of +10 ms, and were repeated 2-70 times at a frequency of 0.5 Hz to elicit t-LTP. In some experiments, longer time windows (positive spike timings: Δ t= +17-25 ms (binned as 20 ms data), Δ t= +38-43 ms (binned as 40 ms data), and Δ t= +96-103 ms (binned as 100 ms data) were used to test for t-LTP. Likewise, short negative spike timings (Δ t= -15ms) were used to test effects of “anti-causal” synaptic stimulation. EPSPs were monitored every 20 s (i.e., 0.05 Hz) for 10 min baseline and then 30 min or 60 min after STDP induction. Unpaired stimulation of 4 postsynaptic APs instead of a full STDP protocol were performed (i.e., 6x 0:4) in a subset of cells that served as controls. In another set of cells, we assessed possible spontaneous changes in synaptic transmission (stimulation at 0.05 Hz for 40 min) in the absence of any STDP stimulation. These recordings served as negative controls (designated 0:0 controls).

To investigate whether a rise of postsynaptic Ca^2+^ concentration is required for induction of t-LTP under our conditions, we applied the Ca^2+^ chelator BAPTA (10 mM, Sigma, Germany) via the patch pipette solution into the recorded postsynaptic neuron. NMDA receptor (R) dependency was tested by application of an NMDAR antagonist (APV 50 µM, DL-2-Amino-5-phosphonopentanoic acid, Tocris, Germany) in the bath solution. The contribution of L-type voltage gated Ca^2+^ channel activation to t-LTP was evaluated with bath applied Nifedipine (25 μM, Sigma, Germany). To interfere with group I metabotropic glutamate receptor (mGluR) signaling we used bath application of either the mGlu_1_ receptor antagonist YM298198 (1 μM, Tocris, Germany) or the mGluR_5_ receptor antagonist MPEP (10 μM, Tocris, Germany) alone, or both blocker simultaneously (the substances were bath applied for a minimum of 15 min prior and during STDP recordings). IP3 receptors were blocked by bath application of 2-APB (100 µM, Tocris, Germany, micro-perfusion pump), 2-APB was applied at least 15 min prior t-LTP induction and was present throughout the recordings. Intracellular infusion of ryanodine (100 µM, Tocris, Germany, infusion for 15 min) was used to block ryanodine receptors of internal calcium stores. Where appropriate, respective controls were performed with ACSF or internal solution containing the same final concentration of DMSO as used for the drug containing solution (i.e. solvent controls) using the same perfusion conditions.

We investigated dopaminergic neuromodulation of STDP by bath application of specific antagonists for D1-like (SCH23390, SCH; 10 μM, Sigma) and D2-like dopamine receptors (Sulpiride, Sulp; 10 μM, Sigma, substances were applied for at least 15 min prior STDP recordings). The contribution of BDNF/TrkB signaling was tested by bath application of a scavenger of endogenous BDNF (recombinant human TrkB-Fc chimera, R&D Systems, Germany). For scavenging of BDNF, slices were pre-incubated for at least 3 h with 5 μg/ml TrkB-Fc, and subsequent recordings were performed in the presence of 100 ng/ml TrkB-Fc (compare Edelmann et al., 2015). Positive controls were recorded in slices kept under the same regime, but without the addition of TrkB-Fc. To test low repeat t-LTP under conditions of chronic 50% BDNF reduction, we used heterozygous BDNF^+/-^ mice and respective wildtype littermates as described previously (Edelmann et al., 2015).

The contribution of activity-dependent incorporation of GluA1 subunit containing AMPA receptors to expression of low repeat t-LTP was verified by postsynaptic application of Pep1-TGL (100 µM, Tocris, Germany) via the patch pipette solution. To investigate a possible role of GluA2 lacking calcium permeable (cp-) AMPA receptors, we used bath applied NASPM (1-Naphtyl acetyl spermine trihydrochloride, 100 µM, Tocris, Germany) or IEM-1460 (*N,N,H*,-Trimethyl-5-[(tricyclo[3.3.1.13,7]dec-1-ylmethyl)amino]-1-pentanaminiumbromide hydrobromide, 100 µM, Tocris, Germany). To test for NO and endocannabinoids as possible retrograde messengers of presynaptically expressed 6x 1:1 t-LTP, we added the CB1 receptor antagonist AM-251 (3 µM; Tocris, UK), or the NO synthase inhibitor L-NAME (200 µM; Tocris, UK) to our recording ACSF.

### Data acquisition and Data Analysis

Data were filtered at 3 kHz using a patch clamp amplifier (EPC-8, HEKA, Germany) connected to a LiH8+8 interface and digitized at 10 kHz using PATCHMASTER software (HEKA, Germany). Data analysis was performed with Fitmaster software (HEKA, Germany). All experiments were performed in the current clamp mode, except for paired pulse ratio (PPR) and miniature EPSCs that were recorded in voltage clamp mode at -70 mV holding potential, as well as AMPA/NMDA receptor current ratios that were recorded in voltage clamp at -70 mV and -20 mV holding potential. The holding potential for recording of NMDAR currents was set to the maximal depolarized value (i.e. -20 mV) that allowed stable recordings in spite of activated voltage gated K^+^ currents. We did not replace K^+^ for Cs^+^ in our internal solutions, since we wanted to elicit LTP under conditions of physiological ion composition and AMPAR/NMDAR current ratio had to be measured before and 30 min after t-LTP induction. Series resistance of cells at start of recordings ranged from 8-30 M**Ω**. Input resistance was continuously monitored by hyperpolarizing current steps (250 ms; 20 pA), elicited prior to evoked EPSP responses. The average EPSP slope calculated from 10 min control recording (baseline) was set to 100% and all subsequent EPSP slopes of a cell were expressed as percentage of baseline slopes. Synaptic potentiation was calculated from the mean EPSP slopes 20-30 min (or 50-60 min) after STDP induction, divided by the mean EPSP slope measured during 10 min before STDP stimulation (baseline). Spike timing intervals (i.e. *Δt*, ms) were measured as the time between onset of the evoked EPSP and the peak of the first action potential. Cells were only included for analysis if the initial resting membrane potential (RMP) was between -55 and -70 mV. Cells were excluded when series resistance was >30 MΩ or when input resistance varied more than 25% over the entire experiment. Furthermore, traces showing visible “run-up” or “run-down” during baseline recording were excluded. In graphs, data were binned at 1 min intervals.

AMPA/NMDA receptor mediated current ratios were calculated from the peak current amplitudes of the fast AMPA receptor mediated components evoked at a holding potential of -70 mV divided by the amplitudes of the NMDAR mediated slow current components measured 50 ms after the onset of EPSCs at a holding potential of -20 mV. Selectivity of this procedure for AMPAR and NMDAR mediated currents was confirmed by bath application of either 50 µM APV or 10 µM NBQX in selected experiments (compare Edelmann et al., 2015).

For analysis of presynaptic short-term plasticity before and after t-LTP induction, paired pulse facilitation was recorded in voltage clamp mode at a holding potential of -70 mV, and the paired pulse ratio (PPR) was determined by dividing the peak current amplitudes of the second EPSC by the amplitude of the first EPSC at an inter-stimulus interval of 50 ms. For cells showing successful t-LTP (i.e. >105% of baseline), the magnitude of PPF before t-LTP induction (−10 min) and after t-LTP expression (+30 min) was plotted as in-cell comparisons in **Fig. 3A**). A decrease in PPF after vs. before t-LTP expression was considered to indicate enhanced transmitter release and thereby revealing a presynaptic contribution to the observed t-LTP.

As an additional measure for possible t-LTP induced presynaptic changes, miniature excitatory postsynaptic currents (mEPSCs) were collected from 3 min of continuous recordings before and 30 min after t-LTP induction. However, while mEPSC recording is usually performed in the presence of TTX, we had to omit TTX here, since t-LTP induction required action potential firing. Thus, we analyzed with the Minianalysis program (Synaptosoft, USA) all small amplitude EPSCs (cut-off amplitude: 20 pA to minimize collection of AP-driven EPSCs), and named these events *putative* mEPSCs in the manuscript. The cut-off frequency was determined from analysis of mEPSCs recorded in CA1 neurons under identical experimental conditions as in t-LTP experiments, but in the presence of TTX (>99% of mEPSC were <20 pA; compare **Fig. S3**). For cells showing successful t-LTP (i.e. >105% of baseline), cumulative fraction plots for inter-event intervals (IEI) of putative mEPSCs were generated. In addition, mean amplitudes of putative mEPSCs before vs. after successful induction of t-LTP, respectively, were compared. To verify independent expression of the two low repeat paradigms, we performed an occlusion approach. Here, we subsequently induced first 6x 1:1 t-LTP and 25 min later 6x 1:4 t-LTP in the same cell, and compared the change in synaptic strength by the two protocols.

To assure reproducibility of results, data for experiments shown in **Figs. 1, 3, 5, 9, and S1** were pooled from 2 or 3 independent experimenters, blind to the results of the other(s).

**Figure 1:**
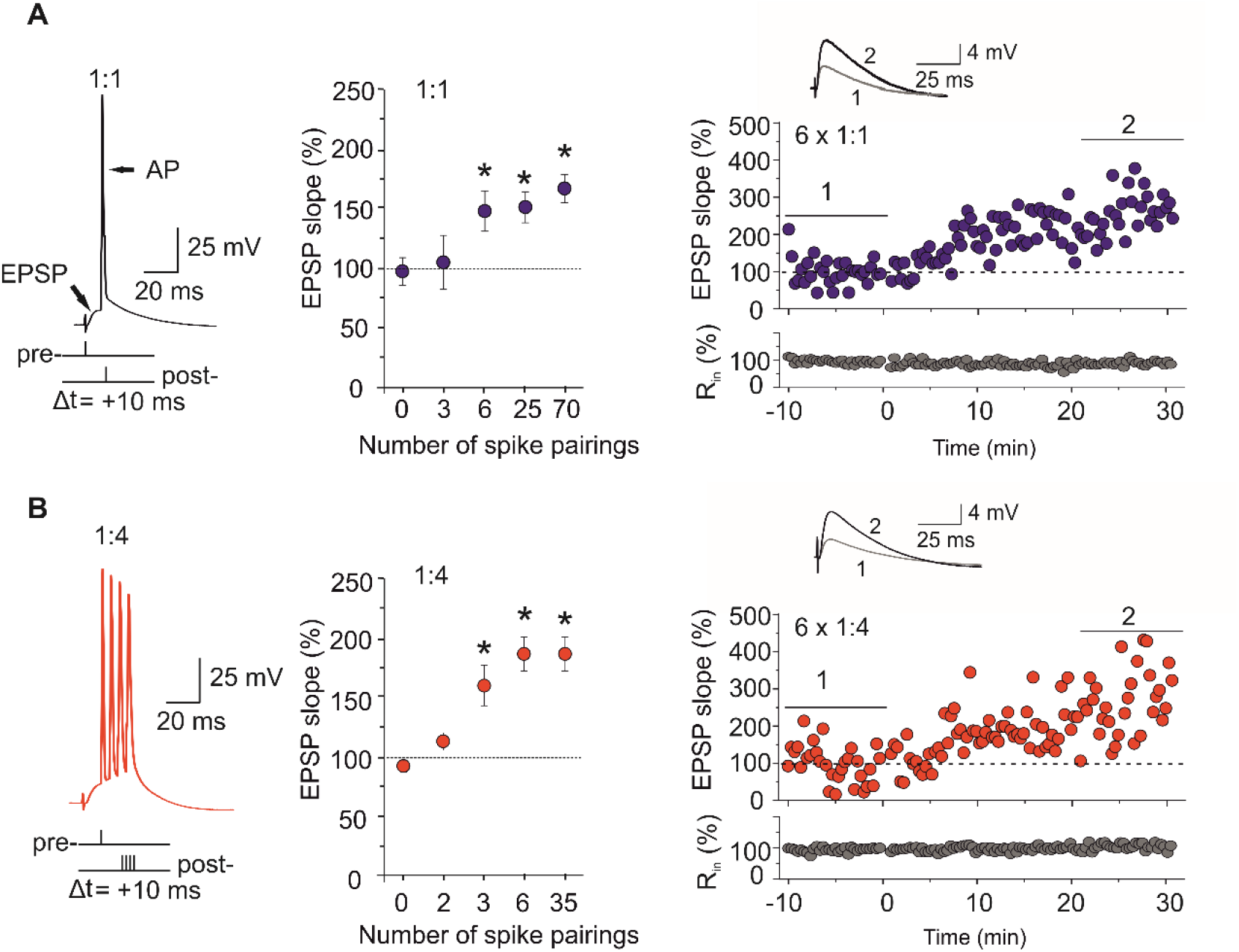
Low repeat t-LTP at Schaffer collateral-CA1 synapses is dependent on stimulation pattern and on the repeat number. **A)** 1:1 t-LTP protocol consisting of the temporal coincident stimulation of an EPSP and one postsynaptic action potential (AP) with a time delay of +10 ms. Summary plot showing the average magnitude of t-LTP (EPSP slope normalized to baseline recording) in response to variable numbers of repeats of the 1:1 t-LTP protocol delivered at 0.5 Hz. Successful t-LTP could be induced with 6 (n=10 / N=7), 25 (n=10 / N=4) and 70 (n=10 / N=7) repeats of the 1:1 t-LTP protocol. Three repeats did not cause any changes in synaptic strength (n=7 / N=6), and showed similar normalized EPSP slopes as negative controls (0:0; n=10 / N=8). Right: original EPSPs and time course of 6x 1:1 induced potentiation recorded in an individual representative cell. **B)** 1:4 t-LTP protocol consisting of the temporal coincident stimulation of an EPSP and four postsynaptic action potentials (APs) with a time delay of +10 ms. Summary plot showing the average magnitude of t-LTP (EPSP slope normalized to baseline recording) in response to variable numbers of repeats of the 1:4 t-LTP protocol delivered at 0.5 Hz. Successful t-LTP could be induced with 3 (n=10 /N= 3), 6 (n=10 / N=6) and 35 (n=10 / N=8) repeats of the 1:4 t-LTP protocol. Two repeats did not cause changes in synaptic strength (n=10 /N =3) and showed similar normalized EPSP slopes as negative controls (0:0; n=7 / N=3). Right: original EPSPs and time course of 6x 1:4 induced potentiation recorded in an individual representative cell. Averaged traces for EPSPs before and after LTP induction are shown as insets. * p< 0.05 ANOVA posthoc Dunnet’s test. Data are shown as mean ± SEM.

### Statistics

Statistical analysis was performed using GraphPad Prism version 8.4 (GraphPad Software, USA) or JMP 8 (SAS Institute Inc., USA). Pooled data of experiments from at least three different animals are expressed as mean ± SEM. The Shapiro Wilk-Test was used to test for normal distribution of data. Paired and unpaired two tailed Student’s t-tests were used for data with normal distribution. Otherwise, the nonparametric Mann-Whitney U*-*test was applied. Multiple comparisons were assessed with a one-way analysis of variance (ANOVA), followed by a post hoc t-test (Dunnet’s test, or Kruskal-Wallis test), or followed by post hoc Dunn’s test for parametric and nonparametric data, respectively. A p-value <0.05 was set as level of significance and is indicated by an asterisk. The actual statistic procedures used for each experiment are mentioned in the text. The respective number of experiments (n) and the number of animals (N) is reported in the figure legends.

## Results

### Timing-dependent LTP at Schaffer collateral-CA1 synapses requires 3-6 spike pairings

Using whole cell patch clamp recordings, we investigated timing-dependent (t-)LTP at Schaffer collateral (SC)-CA1 synapses in acute hippocampal slices obtained from juvenile (i.e. 1 month old) male C57BL/6J mice. T-LTP was induced by STDP protocols consisting of low repeat coincident pre- and postsynaptic action potentials (APs) at low pairing frequencies (0.5 Hz). To limit the number of stimulated SC axons we used glass pipettes (10-15 µm diameter) filled with saline as extracellular stimulation electrodes (compare Methods). Recorded mean EPSP amplitudes before t-LTP induction were ≤ 7 mV. Also, to avoid synaptic network activity by spontaneous AP firing of CA3 neurons, all SCs were cut in CA2 (see Methods). All these precautions were used to assure that the recorded CA1 neuron did not receive uncontrolled evoked excitatory synaptic input. Accordingly, we observed neither multiple component nor polysynaptic EPSPs in the recorded CA1 neuron in response to our extracellular stimulation, nor spontaneously occurring synaptic network activity in the interval between two successive stimulations. SC-CA1 synapses were repeatedly activated by pairing of an excitatory postsynaptic potential (EPSP) that was elicited by supra-threshold extracellular SC stimulation, with a single postsynaptically evoked AP (1EPSP/1AP or 1:1, Δ t= +10 ms at 0.5 Hz; compare **Fig. 1A**). To determine the minimal repeat number required for successful t-LTP induction, neurons were subjected to either 70, 25, 6, or 3 repeats of single spike pairings. Unexpectedly, we found that just six spike pairings with 1:1 stimulation delivered at a frequency of 0.5 Hz were sufficient to induce robust potentiation of EPSP slopes to 147.9 ± 17.0% at 30 min after induction. The t-LTP magnitude was similar to the respective magnitude of t-LTP induced with either 25 or 70 repeats and significantly different from negative controls (25x: 151.1 ± 13.0% and 70x: 166.9 ± 11.8%; ANOVA F_(4,42)_=4,2387 p=0.0057). STDP experiments performed with 3x 1:1 stimulation at 0.5 Hz, showed only a very slight average increase of EPSP slopes to 104.6 ± 22.5% 30 min after induction that was highly variable between cells. The average value was not significantly different from the respective EPSP slopes observed after 40 min (i.e. 10 min baseline + 30 min test) in control neurons that were not subjected to STDP stimulation (negative controls (0:0): 97.2 ± 11.5%; **Fig. 1A**).

The time course of changes in synaptic strength in an individual cell that was potentiated with the low repeat 1:1 protocol is depicted at the right side in blue. These data indicate that t-LTP can be induced at SC-CA1 synapses with low repeat t-LTP paradigms (i.e. 6x 1:1) that might more closely resemble the natural pattern of pre- and postsynaptic activity that can be observed during memory formation in CA1 *in vivo* than any high frequency LTP or high repeat t-LTP protocol.

Since it had been suggested that successful induction of t-LTP at SC-CA1 synapses requires firing of multiple postsynaptic APs (Buchanan and Mellor, 2007; Pike et al., 1999; Remy and Spruston, 2007), we incorporated a postsynaptic burst (4 APs delivered at 200 Hz) into the 6x 1:1 protocol (**Fig. 1B**). This protocol is referred to as 6x 1EPSP/4AP (or 6x 1:4, **Fig. 1B**) paradigm and induced t-LTP at SC-CA1 synapses with the same efficiency as the 6x 1:1 protocol (compare **Fig. 2C, D**). As for the canonical protocol, we determined the threshold number of repeats also for the burst protocol.

**Figure 2:**
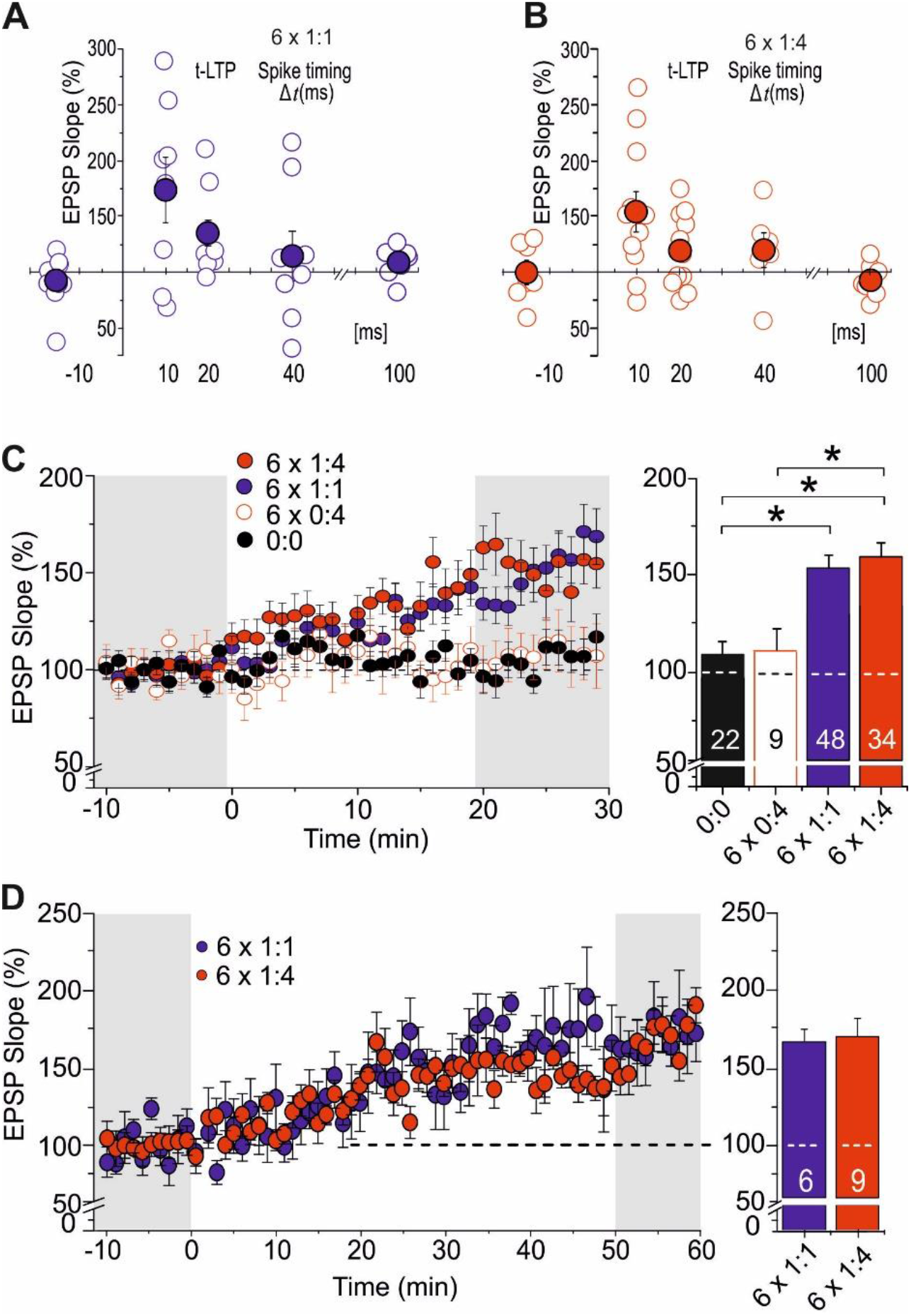
Comparison of canonical and burst low repeat STDP paradigms. STDP plots for 6x 1:1 (**A**, blue) and 6x 1:4 (**B**, red) protocols. Changes in synaptic strength are shown for different intervals between start of the EPSP and postsynaptic APs. Groups of cells at +10 ms interval (standard), with longer time interval (+20, +40 and +100 ms) or with negative spike pairings (−15 ms) are shown. Each open circle represents the result for an individual CA1 neuron. Mean values are shown as closed circles. **C)** The time courses of t-LTP expression did not differ between 6x 1:1 (n= 48 / N= 44) and 6x 1:4 (n=34 / N=27) paradigms, but they were significantly different from negative controls (0:0; n=22 / N=17) and unpaired controls (6x 0:4; n=9 / N=7). The average magnitude of t-LTP is shown in the bar graphs. **D)** Extended measurements for 1 hour after t-LTP induction for both low repeat STDP protocols (6x 1:1: n=6 / N=6, 6x 1:4: n=9 / N=8). Data are shown as mean ± SEM, scale bars are shown in the insets.

As shown in Figure 1B, successful 1:4 t-LTP could be induced by only three repeats of our burst protocol. Three repeats (159.4 ± 15.9%), 6 repeats (185.9 ± 14.2%) and 35 repeats (138.0 ± 7.1%) of 1:4 stimulation all yielded significant potentiation compared to the negative control (0x repeats (92.2 ± 5.0%); ANOVA F_(4,42)_=9.3654 p<0.0001, posthoc Dunnett’s Test: 3x: p=0.0017; 6x: p<0.0001; 35x: p=0.0415). The time course of change in synaptic strength of a typical cell obtained with 6x 1:4 t-LTP stimulation is shown on the right.

Since post-tetanic potentiation is lacking under these stimulation conditions, we observed for both t-LTP protocols a delayed onset (∼5 min) and a subsequent gradual increase of t-LTP magnitude that typically proceeded until 30 min after induction, being consistent with previous t-LTP studies (compare e.g., Banerjee et al., 2009; Edelmann et al., 2015; Edelmann and Lessmann, 2011; Meredith et al., 2003; Nevian and Sakmann, 2006; Pattwell et al., 2012).

For further comparison and analysis of signaling and expression mechanisms of t-LTP we focused in all subsequent experiments on the 6 repeat protocols that induced with the same number of repeats successful t-LTP for both, the canonical and the burst paradigm (indicated as 6x 1:1 or 6x 1:4).

### Canonical and burst containing low-repeat STDP paradigms induce Hebbian t-LTP

When spike pairings were delivered with longer time delays between pre- and postsynaptic firing (Δt: +20, +40, +100 ms) the magnitude of t-LTP declined (Δt: +20 ms: 6x 1:1= 132.1 ± 15.4%, and 6x 1:4= 118.3 ± 9.9%; Δt: +40 ms: 6x 1:1= 116.0 ± 23.2%, and 6x 1:4= 119.19 ± 14.81%; Δt: +100 ms 6x 1:1= 109.0 ± 5.7%, and 6x 1:4= 92.2 ± 5.5%; compared to 6x 1:1= 174.0 ± 29.5% and 6x 1:4= 154.4 ± 18.1% at Δt: +10 ms). Stimulation with short negative time delays (post-pre; Δt: -15 ms) did neither induce t-LTP nor significant t-LTD (6x 1:1: 86.9% ± 9.6%; 6x 1:4: 100.3 ± 9.8%; **Fig. 2A, B**). ANOVA revealed a significant effect of spike timing for 6x 1:1 (F_(4,33)_=32.8186, p=0.0408) and 6x 1:4 t-LTP (F_(4,37)_=3.3328, p= 0.0199). Posthoc Dunnett’s tests revealed significant differences for most comparisons (for 6x 1:1: +10 vs. -15 ms; +10 vs. +40 ms; +10 vs +100 ms; for 6x 1:4: +10 vs. -15 ms; +10 vs. +20 ms; +10 vs. +100 ms). Together these findings indicate that both low repeat t-LTP protocols show the typical characteristics of STDP.

In general, the magnitude of t-LTP induced with the 6x 1:4 stimulation (159.6 ± 8.0%) was comparable to that observed for 6x 1:1 stimulation (153.7 ± 8.2%; p > 0.05, compare **Fig. 2C**). Eliciting only postsynaptic bursts without pairing to presynaptic stimulation (6x 0:4; 110.1 ± 11.9%) did not yield significant potentiation compared to negative controls (0:0; 108.6 ± 7.1%, p > 0.05; **Fig. 2C**). Importantly, these results demonstrate that repeated postsynaptic burst firing alone does not induce any change in synaptic strength, indicating hebbian features for our low repeat t-LTP paradigms, thereby delimiting our protocols from non-hebbian behavioral time scale synaptic plasticity recently reported for hippocampal places cells (Bittner et al., 2017).

Prolonged patch clamp recordings carried out for 1 hour after pairing showed that both low repeat STDP paradigms yielded comparable t-LTP magnitudes at 60 min as observed 30 min after pairing. These data demonstrate that both low-repeat protocols enable longer lasting changes in synaptic transmission without any decline in magnitude. Moreover, the overall time course of the potentiation was indistinguishable for both types of low repeat t-LTP (**Fig. 2D**, potentiation after 1h: 6x 1:1: 164.0 ± 9.5% and for 6x 1:4: 168.0 ± 14.8%, t_(13)_=-0.20435 p=0.84155). Together, these findings indicate that the low repeat STDP paradigms identified in this study induce Hebbian plasticity selectively at short positive spike timings with similar properties as have been described in earlier studies using high repeat canonical and burst type STDP protocols (e.g., Bi and Poo, 1998; Edelmann et al., 2015; Froemke et al., 2006).

In the next series of experiments, we aimed to determine the mechanisms of induction and expression as well as the intracellular signaling cascades involved in modulation of both types of low repeat t-LTP. Of note, the 6x 1:4 protocol (but not the 6x 1:1) was equally potent in inducing t-LTP in the absence or presence of GABA_A_ receptor mediated inhibition (in the absence of picrotoxin: 6x 1:4 t-LTP: 164.38 ± 20.82%, n=10 / N=6, and 6x 1:1 t-LTP: 109.38 ± 13.19%, n=10 / N=5). To assure that all pharmacological treatments in our study affected directly the investigated SC-CA1 glutamatergic synapses rather than mediating their effects through GABAergic inhibitory circuit components, all t-LTP experiments were performed in the presence of GABA_A_ receptor inhibitors (see Methods).

### Influence of single and multiple postsynaptic action potentials on t-LTP expression

To investigate whether the low repeat STDP paradigms introduced here, rely on pre- or postsynaptic expression mechanisms of synaptic plasticity, we determined changes in the paired pulse ratio (PPR) before and 30 min after t-LTP induction that was obtained when two successively evoked EPSPs were elicited at 50 ms inter-stimulus interval (**Fig. 3A**). Commonly, a decrease in PPR after induction of LTP is interpreted as an increase in transmitter release probability and would be expected in case of presynaptically expressed synaptic plasticity. When t-LTP was induced with the 6x 1:1 paradigm, we found on average a significant decrease in PPR (before: 2.15 ± 0.17, after: 1.78 ± 0.12; paired Student’s t-test, t_(14)_ =3.050;p=0.0086. Although we observed also a tendency towards reduced PPR for the 6x 1:4 t-LTP, this change did not reach statistical significance (before: 1.78 ± 0.15; after: 1.56 ± 0.09; paired Student’s t-test, t_(17)_ =1.395; p=0.1811; for negative controls: before: 1.83 ± 0.08, after: 1.93 ± 0.23; paired Student’s t-test, t_(9)_ =0.5128; p=0.6204; **Fig. 3A**). The significantly decreased PPR after induction of 6x 1:1 t-LTP hints at a presynaptic expression mechanism, although the difference to 6x 1:4 t-LTP was modest. Interestingly, the initial PPR before inducing t-LTP was not significantly different between the tested groups (Kruskal-Wallis test, H_(2)_ = 0.8503; p = 0.6537), indicating that the initial release probability was similar and a stable basal parameter in our slices.

**Figure 3:**
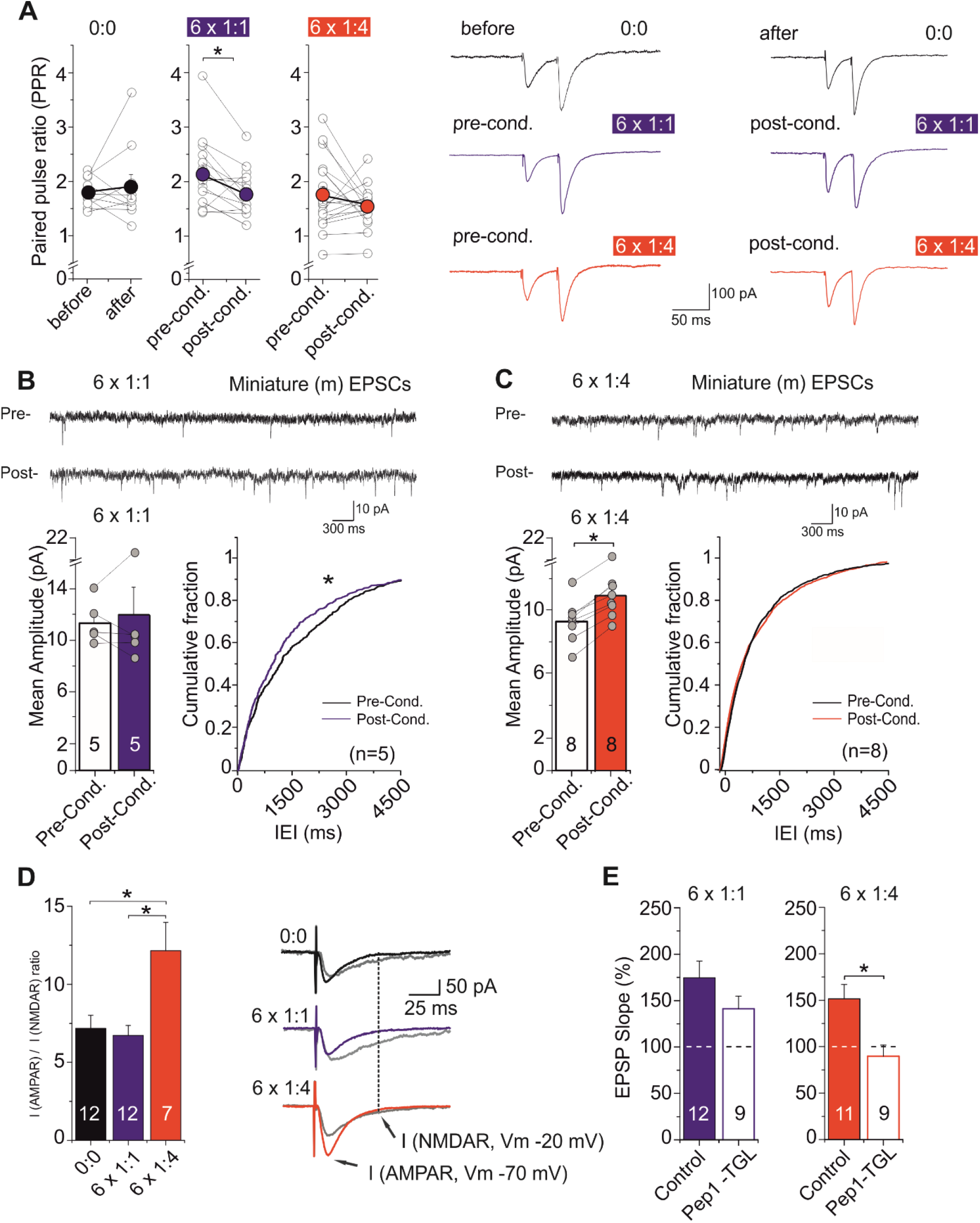
Different loci of expression for t-LTP induced by the low repeat 1:1 and 1:4 paradigms. **A)** Paired pulse ratio (PPR) calculated before (pre-cond.) and 30 min after (post-cond.) t-LTP induction for the same cells (6x 1:1: n=15 / N=6; 6x 1:4: n=18 / N=6), or at the beginning and the end of measurements in negative controls (0:0: n=10 / N=5). **B, C)** Mean amplitudes and cumulative fraction of inter-event intervals (IEI) of putative mEPSCs (see Methods) before and after 6x 1:1 t-LTP induction (**B**), and 6x 1:4 t-LTP protocol (**C**). The significant change in IEI for 6x 1:1 t-LTP in the absence of a significant change in mean amplitudes of putative mEPSCs suggests a presynaptic change in release probability. For 6x 1:4 t-LTP, the significant change in amplitude of putative mEPSC in the absence of change in IEI is consistent with a postsynaptic change. **D)** Right: original traces of AMPAR mediated currents recorded in voltage clamp at -70 mV holding potential (black) and NMDAR (gray) mediated currents at -20 mV holding potential. Left: ratio of AMPA/NMDA receptor mediated currents (AMPAR: peak current at -70 mV; NMDAR: current amplitude 50 ms after start of EPSC, recorded at -20 mV) for negative controls (0:0; n=12 / N= 9) after successful induction of t-LTP with both low repeat paradigms (6x 1:1: n= 12 / N=11; 6x 1:4: n= 7 / N= 5). The increased AMPA/NMDA ratio only after induction of 6x 1:4 t-LTP indicates a postsynaptic change. **E)** Intracellular application of Pep1-TGL (inhibiting membrane insertion of GluA containing AMPARs) via the patch pipette blocked 6x 1:4 t-LTP, whereas 6x 1:1 t-LTP remained intact (6x 1:1: ACSF; n= 12 / N= 10, Pep1-TGL n= 9 / N= 7 and 6x 1:4: ACSF n= 11 / N= 11, Pep-TGL n= 9 / N= 7). Data are shown as mean ± SEM. Scale bars are shown in the figures.

As an additional measure for presynaptic vs. postsynaptic expression mechanism, we determined putative mEPSCs (compare Methods) before and after successful induction of 6x 1:1 and 6x 1:4 t-LTP (Fig.S1, compare Bender et al., 2009; Edelmann et al., 2015). The respective results were consistent with a presynaptic change in release probability for 6x 1:1 t-LTP, as putative mEPSC inter-event intervals (IEI) were significantly decreased after successful 6x 1:1 t-LTP induction (**Fig. 3B**; Kolmogorov-Smirnov 2-sample test: Z=1.5745, p=0.0132), thus indicating a frequency increase of putative mEPSC, whereas amplitudes of putative mEPSC remained unchanged (paired Students t-test, p=0,42; **Fig. 3B**). In contrast, the same analysis revealed for 6x 1:4 t-LTP unchanged frequencies (p=0.32) but significantly enhanced amplitudes of putative mEPSCs (p= 0,017), being consistent with a postsynaptic change in synaptic efficacy.

To further address the locus of expression of 6x 1:1 and 6x 1:4 t-LTP, we analyzed the changes in AMPA/NMDA receptor (R) mediated current ratios 30 min after successful expression of t-LTP. AMPAR mediated peak EPSCs were recorded at a holding potential of -70 mV, while NMDAR mediated current components were determined as remaining current 50 ms after the peak EPSC recorded at -20 mV, to avoid large fluctuations of holding currents that are typically observed at positive membrane potentials. The AMPAR/NMDAR ratio analysis revealed a strong and statistically significant increase in AMPAR-vs. NMDAR-mediated excitatory postsynaptic currents (EPSCs) following successful induction of 6x 1:4 t-LTP, but not when inducing 6x 1:1 t-LTP, or in non-STDP stimulated control cells (0:0: 7.18 ± 0.82, 1:1: 6.72 ± 0.65, 1:4: 12.15 ± 1.81; ANOVA F_(2, 34)_ = 7.979; p = 0.0014, **Fig. 3D**).

Since recording of NMDAR mediated currents at -20 mV (instead of +40 mV) results in smaller current amplitudes we might have introduced a larger error. However, since all groups (negative control, 6x 1:1 and 6x 1:4) were handled identically in this respect, the significant change specifically after t-LTP induction with 6x 1:4 protocol, points to a strong increase of postsynaptic AMPAR conductance, which was absent in the other groups. As an increase in AMPAR/NMDAR mediated currents after inducing LTP is commonly explained by the insertion of new GluA1 containing AMPARs into the postsynaptic spine (Chater and Goda, 2014; Edelmann et al., 2015; Lee and Kirkwood, 2011; Morita et al., 2014) these data strongly suggest a postsynaptic mechanism of expression selectively for the 6x 1:4 t-LTP. Nevertheless, the AMPAR/NMDAR ratio can also be increased by postsynaptic mechanisms other than AMPAR receptor insertion (such as e.g. phosphorylation).

Thus, we next investigated whether the 6x 1:4 t-LTP is mediated specifically by incorporation of GluA1 containing AMPARs. To this aim, we loaded the postsynaptic cells with Pep1-TGL via the patch pipette solution and induced t-LTP with both 6 repeat t-LTP paradigms (compare Edelmann et al., 2015). Pep1-TGL contains the last three amino acids of the C-terminus of the GluA1 subunit, which are required for its insertion into the plasma membrane (Hayashi et al., 2000; Shi et al., 2001). The postsynaptic application of Pep1-TGL resulted in a complete block of 6x 1:4 t-LTP (control: 151.4 ± 15.4%, Pep1-TGL: 89.6 ± 11.9%; Mann-Whitney U test, U = 14.0; p = 0.007). In contrast, t-LTP induced with the 6x 1:1 protocol remained intact under the same recording conditions (control: 174.6 ± 17.9, Pep1-TGL: 141.1 ± 12.1; Mann-Whitney U test, U = 36.0; p =0.201, **Fig. 3E**). Considering these 4 different lines of evidence (PPR analysis, putative mEPSC analysis, AMPA/NMDAR ratio, Pep1-TGL inhibition), our results suggest a dominant postsynaptic locus of expression for the 6x 1:4 t-LTP. In contrast, the absence of any change in AMPA/NMDAR current ratio and AMPAR insertion, in conjunction with the significantly decreased PPR, and the increased frequency of putative mEPSCs indicate a prevailing presynaptic locus of expression for the 6x 1:1 t-LTP. These findings are consistent with our previous results obtained with high repeat t-LTP protocols (Edelmann et al., 2015). The data suggest that the number of postsynaptic spikes fired during induction of low repeat t-LTP decides whether associative Hebbian synaptic plasticity is expressed predominantly by pre- or by postsynaptic mechanisms, whereas the locus of t-LTP expression does not seem to depend on the number of repeats of a specific t-LTP paradigm.

### Distinct calcium sources are recruited for induction of low repeat STDP paradigms

There is a general consensus that induction of long-lasting changes in synaptic strength at SC-CA1 synapses requires a postsynaptic rise in intracellular calcium concentration ([Ca^2+^]_i_) via NMDA receptors (NMDARs, Nicoll and Malenka, 1995). Likewise, also intracellular Ca^2+^ elevation resulting from synchronous activation of NMDARs, L-type voltage-gated Ca^2+^ channels (VGCC), and release of Ca^2+^ from internal stores, following activation of metabotropic glutamate receptors (mGluRs) and subsequent activation of IP_3_ receptors can be responsible for postsynaptic STDP induction (Tigaret et al., 2016).

To verify a role of postsynaptic Ca^2+^ signaling for the induction of 6x 1:4 t-LTP and 6x 1:1 t-LTP, we loaded postsynaptic neurons with 10 mM of the Ca^2+^ chelator BAPTA via the patch pipette solution. After obtaining the whole cell configuration, the BAPTA containing internal solution was allowed to equilibrate for 30 min before t-LTP induction. Likewise, in the respective control experiments t-LTP was also induced 30 min after breaking the seal. As shown in **Fig. 4A and B**, buffering of intracellular Ca^2+^ signals with BAPTA resulted in a complete block of both 6x 1:4 t-LTP (Control: 150.28 ± 12.07%, BAPTA_i_: 104.80 ± 5.84%; Mann-Whitney U test, U = 3.0; p = 0.0303) and 6x 1:1 t-LTP (control: 149.44 ± 15.92%, BAPTA: 103.87 ± 10.37%; unpaired Student’s t-test, t_(10)_= 2.399; p= 0.0374), indicating that a rise in postsynaptic [Ca^2+^]_i_ is required for induction of both low repeat t-LTP forms.

**Figure 4:**
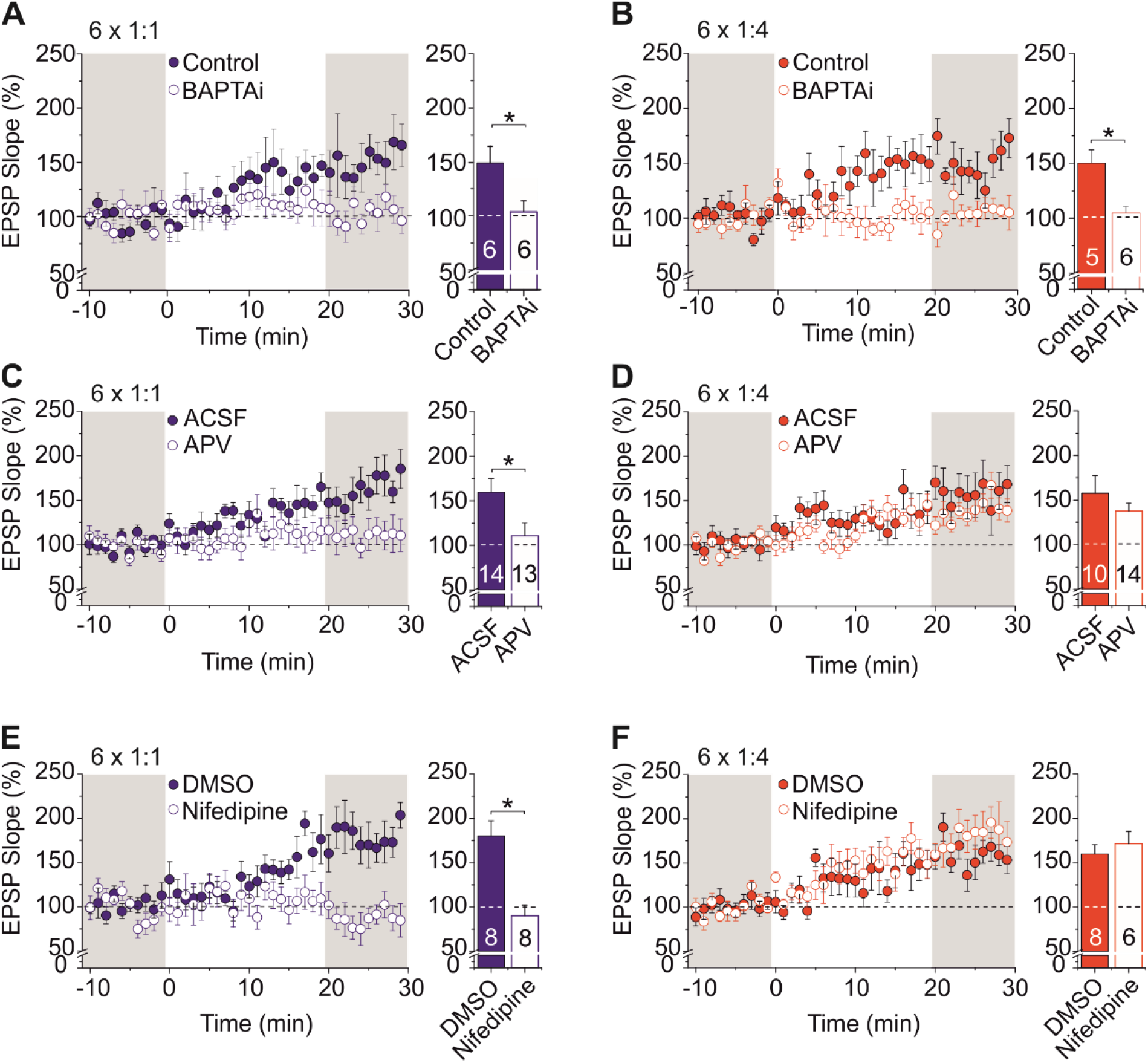
Contribution of NMDA receptors and VGCCs to the induction of 6x 1:1 and 6x 1:4 t-LTP. **A, B)** Inclusion of 10 mM BAPTA in the pipette solution and equilibration with the cell interior for 30 min before t-LTP induction (open circles) prevented t-LTP induced by 6x 1:1 and 6x 1:4 stimulation compared to identically treated (i.e. t-LTP induction 30 min after breaking the patch) control cells (closed circles; Control 6x 1:1: n=6 / N=2, BAPTAi 6x 1:1 : n=6 / N=3; Control 6x 1:4: n=5 / N=5, BAPTA_i_ 6x 1:4: n=6 / N=4), indicating the necessity of postsynaptic calcium elevation to induce t-LTP. **C-F)** Effects of bath applied NMDAR antagonist APV (50 µM) and L-type VGCC inhibitor Nifedipine (25 µM) on low repeat t-LTP. Inhibition of either NMDARs **(C)** or L-type VGCCs **<u>(E)</u>** completely blocked 6x 1:1 t-LTP (**C**: 6x 1:1: ACSF n=14 / N=8, APV n= 13 / N= 12; **E**: DMSO control n= 8 / N=7, Nifedipine n=8 / N=7). **(D)** 6x 1:4 t-LTP remained unaffected by application of the NMDAR inhibitor APV (6x 1:4: ACSF n=10 / N=6, APV n=14 / N=9). **F)** 6x 1:4 t-LTP was not inhibited in the presence of the L-type VGCC inhibitor nifedipine (DMSO n=8 / N=4, Nifedipine n=6 / N=4). Average time course of potentiation and mean (± SEM) magnitude of t-LTP are shown for the respective experiments.

Next, we investigated the sources for the intracellular Ca^2+^ elevation triggering the 6x 1:1 and 1:4 t-LTP. Interestingly, the 6x 1:1 t-LTP was significantly impaired when it was executed either in the presence of the specific NMDAR antagonist APV (50 µM; Control: 159.73 ± 15.23%, APV: 110.91 ± 14.22%; unpaired Student’s t-test, t_(26)_= 2.348; p= 0.0268 ; **Fig. 4C**), or in the presence of the L-type VGCC inhibitor Nifedipine (25 μM; DMSO: 180.11 ± 17.32%, Nifedipine: 90.19 ± 12.15%; unpaired Student’s t-test, t_(14)_= 4.25; p = 0.0008; **Fig. 4E**). In contrast, neither APV (50 μM, unpaired Student’s t-test, t_(22)_ = 1.016; p = 0.3207; **Fig. 4D**) nor Nifedipine (25 μM, Mann-Whitney U test, U= 20.0; p= 0.6620, **Fig. 4F**) inhibited t-LTP induced with the 6x 1:4 protocol (Control: 157.55 ± 19.71%, APV: 137.95 ± 8.39% and DMSO: 159.75 ±10.81%, Nifedipine: 171.73 ± 13.62%). These data demonstrate that postsynaptic Ca^2+^ influx via NMDARs and L-type VGCCs is required for 6x 1:1 t-LTP but not for 6x 1:4 t-LTP induction.

Since induction of t-LTP involves repeated glutamate release that, according to hebbian rules, should contribute to the induction process, we next tested the involvement of metabotropic glutamate receptors (mGluRs) in 6x 1:4 t-LTP. In the hippocampal CA1 region mGluR_1_ and mGluR_5_ are widely expressed and have been reported to induce Ca^2+^ release from internal calcium stores during LTP (e.g., Balschun et al., 1999; Neyman and Manahan-Vaughan, 2008; Wang et al., 2016). Nevertheless, blocking mGluR activation by bath application of antagonists of either mGluR_1_ (YM-298198 10 μM) or mGluR_5_ (MPEP, 10 μM) alone, did not affect the magnitude of 6x 1:4 t-LTP compared to ACSF controls (Control: 159.07 ±17.67%; YM: 155.65 ± 18.70%; MPEP: 143.51 ± 12.43%; Kruskal-Wallis test, H_(2)_= 0.2774; p= 0.8705; **Fig. 5A**). However, co-application of the mGluR_1_ and mGluR_5_ antagonists significantly reduced 6x 1:4 t-LTP magnitude (Control: 178,99 ± 15.29%; YM+MPEP: 129.97 ± 7.94%; unpaired Student’s t-test, t_(22)_ = 2.248; p= 0.0093; **Fig. 5B**), indicating that the activation of one of these receptors alone (either mGluR_1_ or mGluR_5_) is necessary and sufficient to support 6x 1:4 t-LTP.

**Figure 5:**
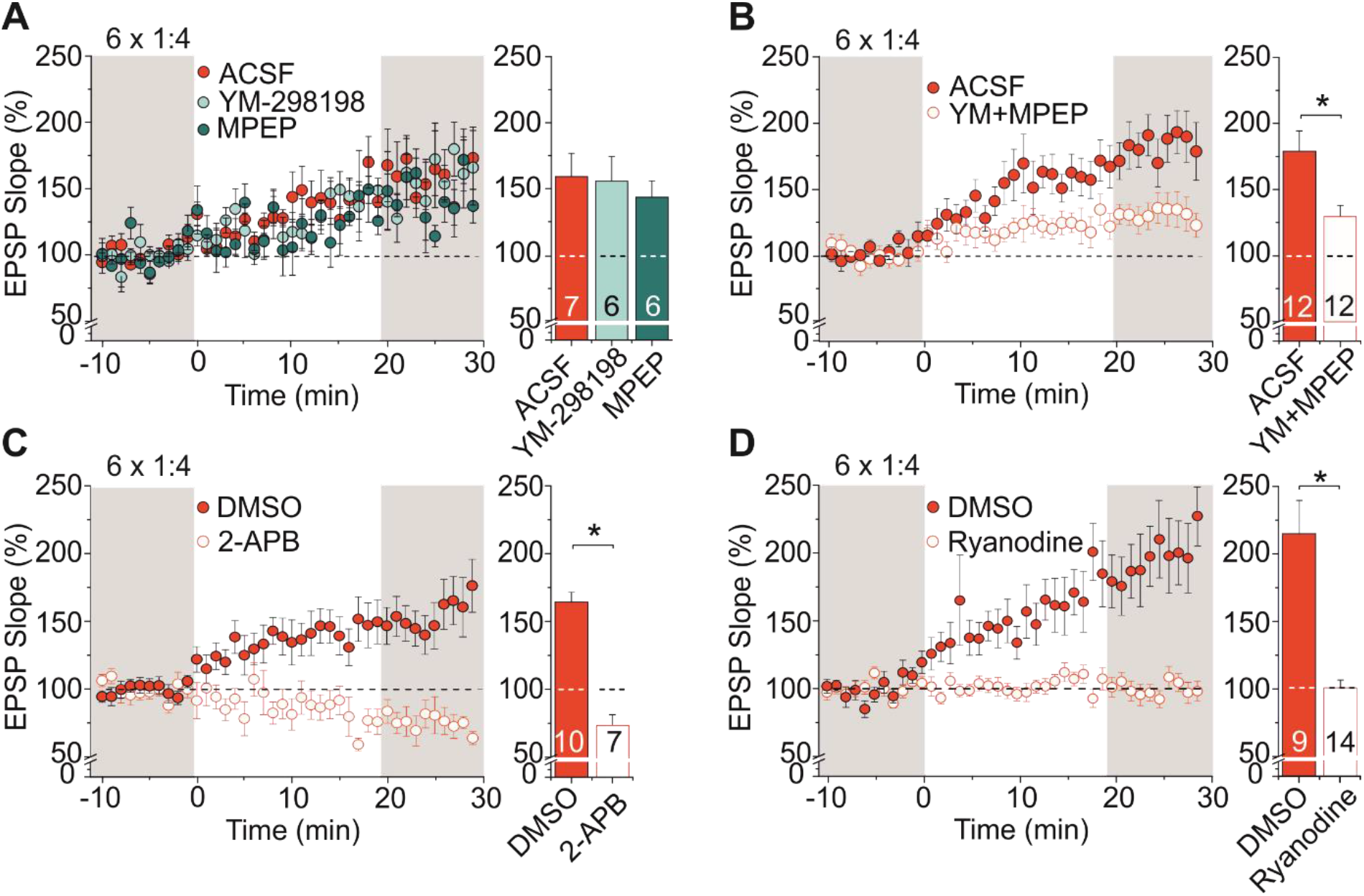
Contribution of group I mGluRs, IP3 receptors and ryanodine receptor-dependent calcium release from internal stores to 6x 1:4 t-LTP. **A)** T-LTP induced with the 6x 1:4 protocol was neither affected by bath application of the mGluR1 antagonist YM-298198 (1 µM; ACSF: n=7 / N=5, YM-298198: n=6 / N=3), nor by the mGluR5 antagonist MPEP (10 µM, n=6 / N=4). **B)** However, co-application of both antagonists (YM-298198 and MPEP; ACSF n=12 / N=8, YM-MPEP n=12 / N=5) significantly reduced synaptic potentiation. **C)** Inhibition of IP3 receptors by 100 µM 2-APB (in 0.05% DMSO) completely blocked 6x 1:4 t-LTP (DMSO n=10/ N=5; 2-APB n=7/ N=3). **D)** Wash in of 100 µM ryanodine into the postsynaptic neuron via the patch pipette inhibited t-LTP induced by 6x 1:4 stimulation (DMSO n= 9 / N= 4; Ryanodine n= 14 / N= 5). Average time course of potentiation and mean (± SEM) magnitude of t-LTP are shown for the respective experiments.

To investigate whether mGluR mediated Ca^2+^ release from internal stores contributes to 6x 1:4 t-LTP we used 2-APB as an inhibitor of IP_3_-receptors. As expected, inhibition of IP_3_-mediated Ca^2+^ release completely blocked 6x 1:4 t-LTP (DMSO: 164.30 ± 18.29; 2-ABP: 71.87± 7.54; unpaired Student’s t-test, t_(15)_=4.0297; p =0.0019, **Fig. 5C**). To examine the involvement of ER-resident ryanodine receptors (RyR) in the low repeat burst protocol, we applied 100 µM ryanodine (a concentration known to irreversibly inhibit RyR; Gao et al., 2005) via the patch pipette into the recorded postsynaptic neurons. As expected in case of RyR involvement, 6x 1:4 t-LTP induction was completely inhibited under these conditions (DMSO: 216.21 ± 25.70%; Ryanodine: 100.24 ± 5.07%; Mann-Whitney U test, U= 5.5; p= 0.0003; **Fig. 5D**).

These data demonstrate that Ca^2+^ release from the ER is a critical component of 6x 1:4 t-LTP. Thus, the postsynaptic Ca^2+^ elevation required for induction of 6x 1:4 t-LTP seems to involve mGluR_1_ or mGluR_5_ mediated release of Ca^2+^ from the ER via IP3 receptors and subsequent Ca^2+^ induced Ca^2+^ release via RyRs (compare **Fig. 10**).

To determine whether internal Ca^2+^ stores also contribute to induction of 6x 1:1 t-LTP, we tested the role of mGluR-, IP3- and RyR-inhibitors in the same way as described above for the 6x 1:4 t-LTP. Interestingly, in contrast to the 6x 1:4 burst protocol, co-application of mGluR1 and mGluR5 antagonists failed to block t-LTP induction by the canonical 6x 1:1 protocol (ACSF: 147.42 ± 11.22%; YM+ MPEP: 145.92 ± 8.55%; unpaired Student’s t-test, t_(11)_= 0.1080; p= 0.9159; **Fig. 6A**). However, the IP3 receptor antagonist 2-APB (DMSO: 139.45 ± 15.37%; 2-ABP: 80.23 ± 8.06%; unpaired Student’s t-test, t_(9)_= 3.590; p= 0.0058, **Fig. 6B**), and the RyR antagonist ryanodine (DMSO: 157.57 ± 11.18% Ryanodine: 117.55 ± 8.35%; unpaired Student’s t-test, t_(17)_= 2.816; p= 0.0119; **Fig. 6C**) both completely blocked 6x 1:1 t-LTP. This suggests that IP3Rs are recruited also by the 6x 1:1 protocol. Since mGluRs are not involved here, Gα_q_/PLC signaling by D1-class dopamine receptors or D1/D2 dopamine receptor heterodimers ((compare Beaulieu and Gainetdinov, 2011)) might be involved (compare **Fig. 10**).

**Figure 6:**
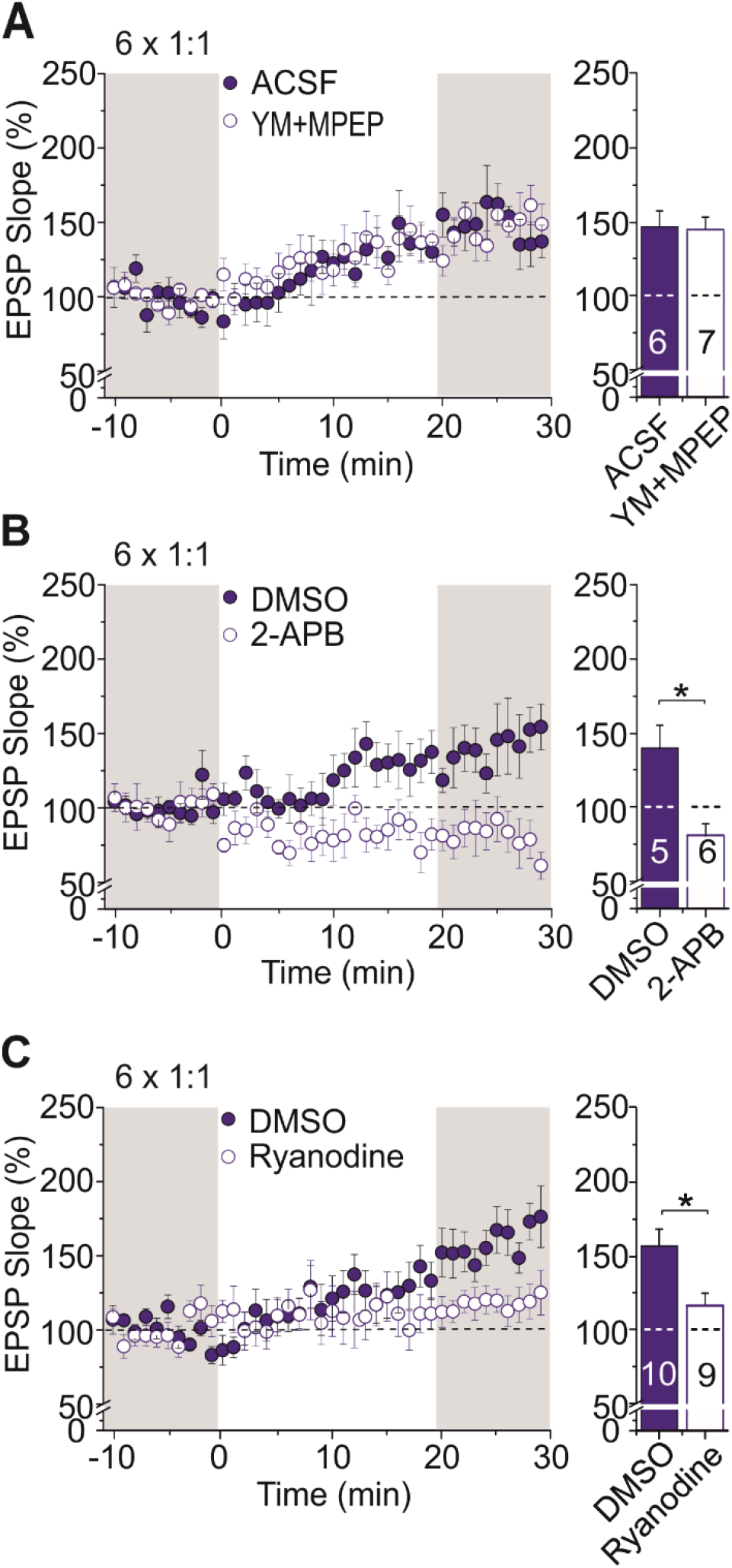
Contribution of group I mGluRs, IP3 receptors and ryanodine receptor-dependent calcium release from internal stores to 6x 1:1 t-LTP. **A)** Co-application of both mGluR antagonists (compare Fig.5; YM-298198 and MPEP; ACSF n=6 / N=3, YM-MPEP n=7 / N=2) did not reduce 6x 1:1 t-LTP. **B)** Inhibition of IP3 receptors by 100 µM 2-APB (in 0.05% DMSO) completely blocked 6x 1:1 t-LTP (DMSO n=5 / N=4; 2-APB; n=6 / N=4). **C)** Wash in of 100 µM ryanodine via the patch pipette into the postsynaptic neuron inhibited t-LTP induced by 6x 1:1 stimulation (DMSO n=10 / N=4, Ryanodine n=9 / N=3). Average time course of potentiation and mean (± SEM) magnitude of t-LTP are shown for the respective experiments.

### Distinct dopaminergic modulation of different low repeat t-LTP protocols at Schaffer collateral-CA1 synapses

Dopamine (DA) serves as an important neuromodulator in learning and memory formation, as well in synaptic plasticity mechanisms underlying both phenomena. DA receptors in the brain are classified into D1-like receptors that include D1 and D5, and D2-like receptors that include D2, D3 and D4 (Missale et al., 1998). It has been shown that activation of D1/D5 receptors has a particularly strong effect on hippocampal synaptic efficacy (Dubovyk and Manahan-Vaughan, 2018; Papaleonidopoulos et al., 2018), and high repeat t-LTP in hippocampal neurons(see e.g.,(Edelmann and Lessmann, 2011; Zhang et al., 2009). Moreover, D4 receptors are known to modulate NMDAR function in CA1 in vivo and ex vivo (Herwerth et al., 2012; Li et al., 2016). Thus, to examine whether in our case, endogenous DA signaling is an essential component of synaptic mechanisms triggering low repeat t-LTP, we investigated the effect of specific bath applied antagonists for D1-like and D2-like DA receptors (D1: SCH23390 (SCH), 10 μM; D2: Sulpiride (Sulp), 10 μM). We found that t-LTP induced with 6x 1:1 stimulation was blocked completely when SCH23390 and Sulpiride were coapplied (Control: 174.15 ± 15.26%, SCH: 165.84 ± 18.03%, Sulp: 136.23 ± 12.99%, SCH+Sulp: 93.02 ± 9.03%; ANOVA F_(3,36)_ = 6.2519; p= 0.0016, posthoc Tukey -test for ACSF vs. SCH+Sulp p= 0.0015 and for SCH vs. SCH+Sulp: p= 0.0091), whereas application of either the D1-like or the D2-like receptor antagonist alone did not significantly reduce the magnitude of the 6x 1:1 t-LTP (**Fig. 7A**). In contrast, the 6x 1:4 t-LTP was dependent exclusively on D2-like receptor signaling, as was evident from complete inhibition of this burst t-LTP in the presence of Sulpiride (significantly different from ACSF controls; Kruskal-Wallis test H _(3)_ = 12.65; p = 0.005, **Fig. 7B**), whereas SCH23390 was without effect (Control: 147.51 ± 8.25%, SCH: 153.64 ± 14.47%, Sulp: 102.34 ± 12.25, SCH+Sulp: 108.67 ± 9.17%).

**Figure 7:**
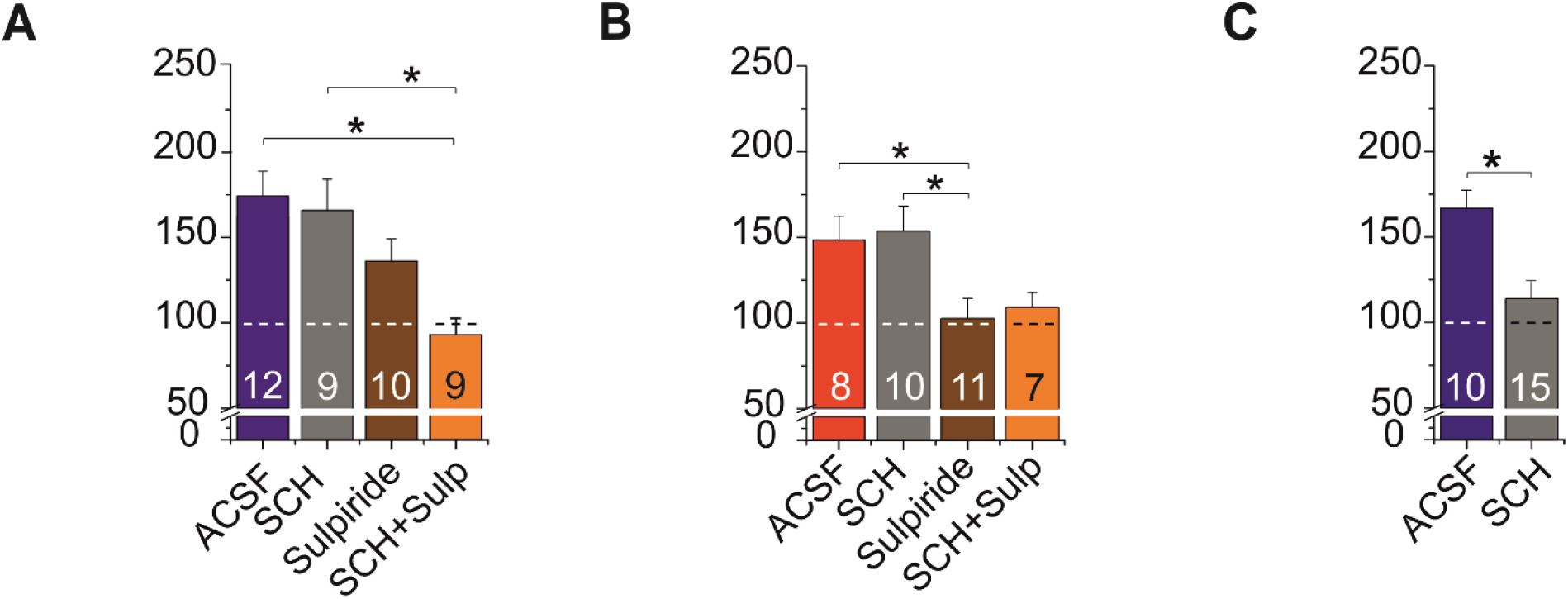
Differential modulation of canonical and burst low repeat t-LTP by dopaminergic signaling. **A)** Dependence of 6x 1:1 t-LTP on D1 and D2 receptor signaling. Neither bath application of SCH23390 (SCH, D1-like antagonist; 10 µM) nor bath application of Sulpiride (Sulp, D2-like antagonist; 10 µM) alone impaired t-LTP (ACSF: n=12 / N=9; SCH23390 n=9 / N=6; Sulpiride n=10 / N=8). However, co-application of both antagonists significantly reduced t-LTP (SCH + Sulp n=9 / N=4). **B)** T-LTP induced with the 6x 1:4 protocol was impaired in the presence of Sulpiride, but not further reduced by co-application with SCH23390. Accordingly, application of SCH23390 alone did not affect 6x 1:4 t-LTP (ACSF: n=8 / N=6; SCH23390 n=10 / N=5; Sulpiride n=11 / N=8; SCH + Sulp n=7 / N=4). **C)** T-LTP induced with the high repeat (70x) 1:1 protocol was inhibited in the presence of the D1 receptor antagonist SCH23390 (10 µM) in mouse slices (ACSF n=10 / N=8; SCH23390 n=15 / N=5) to a similar extent as observed previously in rat hippocampal slices (Edelmann and Lessmann, 2011). Mean (± SEM) magnitude of t-LTP is shown for the respective experiments.

Together, these data indicate that 6x 1:1 t-LTP depends on D1/D2 receptor co-signaling whereas 6x 1:4 t-LTP is only dependent on D2 receptors, highlighting a novel and important role of D2 receptors in both types of t-LTP. This is at variance with the fact that most previous studies investigating DA-dependent conventional LTP at SC-CA1 synapses reported an eminent role of D1-like receptors in high frequency induced LTP forms (e.g., Hagena and Manahan-Vaughan, 2016; Papaleonidopoulos et al., 2018). However, our results are fully consistent with the previously described D2 receptor mediated enhancement of t-LTP in the prefrontal cortex (Xu and Yao, 2010), and the prominent role of D2 receptors in hippocampus-dependent learning (Nyberg et al., 2016). A classical role for D1 receptor signaling was also described for high repeat (70x) canonical t-LTP in rat hippocampal slices (Edelmann and Lessmann, 2011, 2013). To clarify whether repeat number or species matter for the contribution of D1 and D2 receptors in t-LTP, we examined DA dependence of high repeat 70x 1:1 t-LTP in mouse hippocampal slices. We found that also in mouse slices 70x 1:1 t-LTP was fully blocked by bath application of the D1 antagonist SCH23390 (Control: 166.85 ± 11.77, SCH: 113.38 ± 11.07; unpaired Student’s t-test, t _(22)_ = 3.028; p= 0.0062; **Fig. 7C**). These data reveal that high repeat number induced t-LTP is regulated by D1 signaling whereas D2 signaling is selectively involved in low repeat t-LTP. Further, the extent of D2 receptor involvement in low repeat t-LTP is regulated by the postsynaptic spike pattern used for t-LTP induction (compare **Fig. 7A** and **B**).

### The role of BDNF/TrkB signaling in low repeat t-LTP induced by canonical or burst protocols

We recently showed for SC-CA1 synapses that brain-derived neurotrophic factor (BDNF) induced tropomyosin related kinase B (TrkB) signaling mediates t-LTP elicited by a 1:4 t-LTP paradigm with 25 repeats at 0.5 Hz. This t-LTP is driven by an autocrine postsynaptic BDNF/TrkB mechanism that ultimately relies on postsynaptic insertion of new AMPA receptors (Edelmann et al., 2015).

To address whether release of endogenous BDNF might be involved also in low repeat t-LTP, we next tested our low repeat t-LTP protocols in slices obtained from heterozygous BDNF knockout (BDNF^+/-^) mice that express ∼50% of BDNF protein levels compared to WT littermates (e.g., Endres and Lessmann, 2012; Psotta et al., 2015). Our results show that both types of low repeat t-LTP remained functional in response to this chronic depletion of BDNF (6x 1:1 t-LTP: WT: 131.37 ± 9.67, BDNF+/-: 134.47 ± 14.85; Mann-Whitney U test, U = 25.0; p= 0.7789; and 6x 1:4 t-LTP: WT: 134.41 ± 8.78, BDNF+/-: 128.11 ± 8.12; Mann-Whitney U test, U = 14.0; p= 0.5887, **Fig. 8A**).

**Figure 8:**
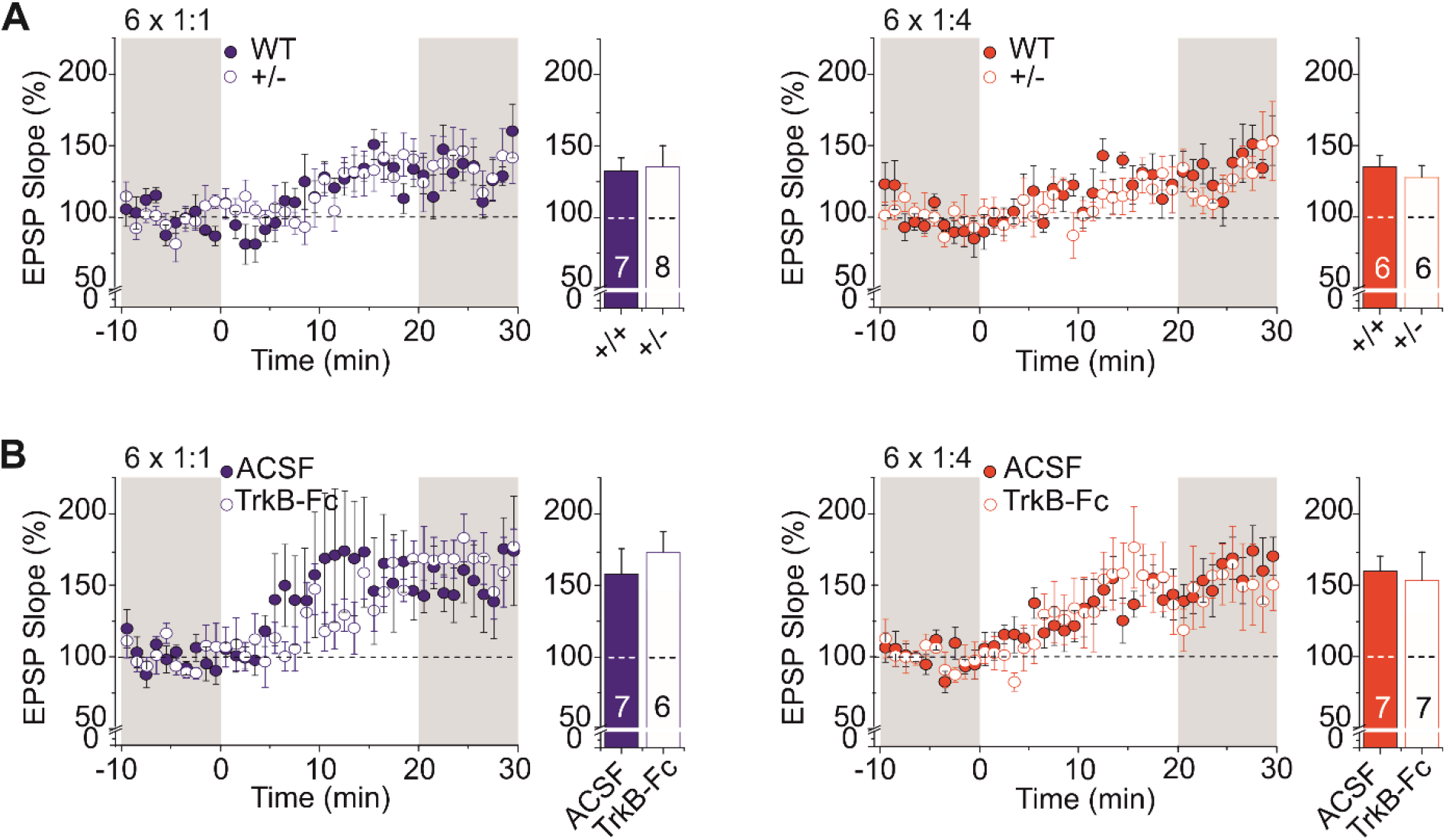
BDNF induced TrkB receptor signaling is not required for t-LTP elicited by low repeat t-LTP protocols. **A)** Low repeat t-LTP was not different for 6x 1:1 (left) and 6x 1:4 (right) stimulation in heterozygous BDNF knockout animals (+/-) compared to wild type litter mates (+/+) (6x 1:1: +/+ n =7 / N=6, +/- n=8 / N=7; 6x 1:4: +/+ n=6 / N=5, +/- n=6 / N=5). **B)** Bath application of the BDNF scavenger TrkB-Fc (100 ng/ml; 3h preincubation) did not affect t-LTP in response to the two low repeat protocols (left: 6x 1:1: ACSF n=7 / N=5, TrkB-Fc n=6 / N=6; right: 6x 1:4: ACSF n=7 / N=5, TrkB-Fc n= 7 / N= 6). Average time course of potentiation and mean (± SEM) magnitude of t-LTP are shown for the respective experiments.

Next, to examine whether acute inhibition of BDNF/TrkB signaling affects low repeat t-LTP, we asked whether scavenging of BDNF by bath applied TrkB receptor bodies (human TrkB-Fc chimera, TrkB-Fc) impairs low repeat canonical or burst t-LTP. However scavenging of BDNF had no effect on the magnitude of t-LTP induced by either of the two protocols (6x 1:1 t-LTP: ASCF: 157.39 ± 18.19, TrkB-Fc: 173.07 ± 14.05; Mann-Whitney U test, U= 13.0; p= 0.0939; and 6x 1:4 t-LTP: ACSF: 159.78 ± 10.36, TrkB-Fc: 152.80 ± 20.28; Mann-Whitney U test, U= 22.0; p= 0.8048, **Fig. 8B**). Together these data indicate that 6x 1:4 and 6x 1:1 t-LTP are both independent from activity-dependent release of endogenous BDNF and downstream TrkB signaling. In conjunction with our previous observation that 25x 1:4 t-LTP is dependent on release of endogenous BDNF (compare Edelmann et al., 2015) the present data suggest that a higher number (>6) of postsynaptic spike bursts in the t-LTP protocol is required to activate BDNF secretion.

### The role of GluA2-lacking, calcium-permeable AMPA receptors in low repeat t-LTP

The transient incorporation of GluA2-lacking, Ca^2+^ permeable (cp-) AMPARs after LTP induction has been proposed as an important process to increase postsynaptic Ca^2+^ levels for LTP expression (Kauer and Malenka, 2006; Man, 2011; reviewed in Park et al., 2018; Plant et al., 2006). To examine whether these receptors are involved in low repeat t-LTP, we incubated our recorded hippocampal slices with the selective cp-AMPAR inhibitor NASPM (100 µM). Interestingly, 6x 1:1 t-LTP and 6x 1:4 t-LTP were both completely blocked in the presence of NASPM (6x 1:1: ACSF: 142.81 ± 12.11, NASPM: 89.45 ± 9.04; unpaired Student’s t-test, t_(14)_ = 3.3502; p= 0.0048; 6x 1:4: ACSF: 177.66 ± 16.83, NASPM: 100.32 ± 5.95; unpaired Student’s t-test, t_(19)_ = 4.829; p= 0.0002, **Fig. 9A, B**). Surprisingly, these results indicate that the influx of Ca^2+^ via GluA2-lacking, cp-AMPARs is mandatory to elicit low-repeat t-LTP induction.

**Figure 9.**
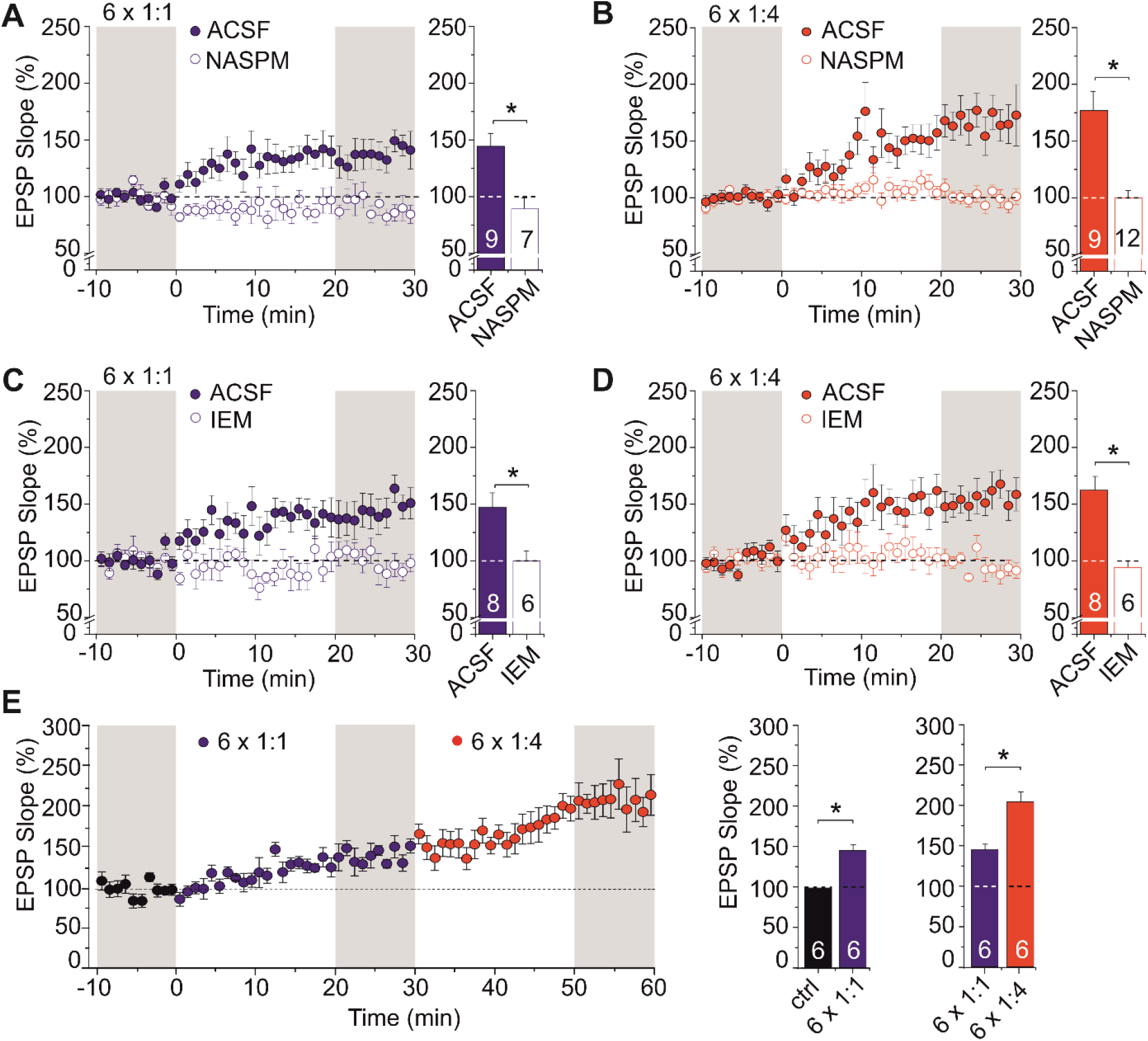
: Ca^2+^ influx via GluA2-lacking calcium-permeable AMPARs is required for low-repeat t-LTP induction. Bath application of a selective inhibitor of Ca^2+^ permeable AMPARs (100 µM NASPM) blocks 6x 1:1 t-LTP (**A**, 6x 1:1: ACSF n=9 / N=5, NASPM n=7 / N=3) as well as 6x 1:4 t-LTP (**B**, 6x 1:4: ACSF n= 9 / N=6 NASPM n=12 / N=5). Also IEM-1460 (100 µM), a second specific inhibitor of cp-AMPARs, blocks 6x 1:1 t-LTP (**C**, 6x 1:1: ACSF n=8 / N=5, IEM-1460 n=6 / N=3) as well as 6x 1:4 t-LTP (**D**, 6x 1:4: ACSF n= 8 / N=5, IEM n=6 / N=3). **E**) Successful induction of 6x 1:1 t-LTP and subsequent 6x 1:4 t-LTP in the same cells (n=6 / N=3). Note the absence of any signs of occlusion between t-LTP induced by the two low repeat protocols (compare **Fig. S2**). Average time course of potentiation and mean (± SEM) magnitude of t-LTP are shown for the respective experiments.

To rule out off-target effects of NASPM, we verified cp-AMPAR contribution in low repeat t-LTP with a second inhibitor of cp-AMPARs (IEM-1460, 100 µM). As shown in **Fig. 9C** and **D**, we observed complete inhibition of low repeat t-LTP also by IEM for both protocols (6x 1:1: ASCF: 146.99 ±12.56, IEM: 99.89 ± 8.43; unpaired Student’s t-test, t_(12)_ = 2.76256; p= 0.0172; 6x 1:4: ASCF: 162.03 ±12.70, IEM: 94.70 ± 6.24; unpaired Student’s t-test, t_(12)_ = 4.4567; p= 0.0007, **Fig. 9C, D**).

In light of the many differences in the induction, expression mechanisms, and dopaminergic modulation of the canonical 6x 1:1 t-LTP and the 6x 1:4 burst t-LTP we asked whether both types of t-LTP can be elicited completely independent from one another or if they occlude each other. To this aim, we first induced 6x 1:1 t-LTP followed in the same cells by a subsequently induced 6x 1:4 t-LTP. As shown in **figure 9E**, both types of t-LTP could be activated independently without any signs of occlusion (1^st^ t-LTP induction (6x 1:1, 143.68 ± 5.85%): t_(5)_=-3.4618; p= 0.0180; 2^nd^ t-LTP induction (6x 1:4, 203.17 ± 12.04%): t_(5)_=-4.7081; p= 0.0053, paired Student’s t-test). Importantly, in another set of cells, subsequent stimulation for a second time with the same 6x 1:1 protocol that had already successfully induced t-LTP, did not yield further potentiation (**Fig. S2**; 1^st^ t-LTP induction (6x 1:1, 147.31 ± 13.06%): t_(4)_=-3.4607; p= 0.0258; 2^nd^ t-LTP-induction (6x 1:1, 157.97 ± 17.88%): t_(4)_=-1.9649; p= 0.1209; paired Student’s t-test). Of note, t-LTP magnitude at 60 min after single 6x 1:1 stimulation amounted to 164.0 ± 9.50% (**compare Fig.2D**), which was not significantly different from t-LTP 60 min after 2 times repeated 6x 1:1 stimulation (157.97 ± 17.88%). This indicates saturated 6x 1:1 t-LTP in response to the first application of the 6x 1:1 protocol.

Given the strong differences in the induction processes and the presynaptic expression of 6x 1:1 vs. postsynaptic expression of 6x 1:4 t-LTP, the absence of occlusion between the two protocols was an expected finding. However, this result highlights the independence of the two different types of low repeat t-LTP investigated here.

The scheme presented in **Fig.10** is consistent with our experimental findings for the prevailing presynaptic expression of 6x 1:1 t-LTP and the predominant postsynaptic expression of 6x 1:4 t-LTP, and depicts the putative roles of mGluRs, cp-AMPARs, dopamine signaling, and internal Ca^2+^ stores in low repeat t-LTP. However, since the distribution of dopaminergic fibers and the pre- and/or postsynaptic dopamine receptor localization in the CA1 region is not yet completely clear (compare Edelmann and Lessmann, 2018), further experiments are clearly required to improve the mechanistic understanding of this aspect of low repeat t-LTP. None withstanding, both low repeat t-LTP forms are already by now clearly distinguishable. Their different features of induction and expression mechanisms and the distinct signaling cascades they employ are likely to form the basis for the versatile computing capacity of individual CA1 neurons in the hippocampus.

**Figure 10:**
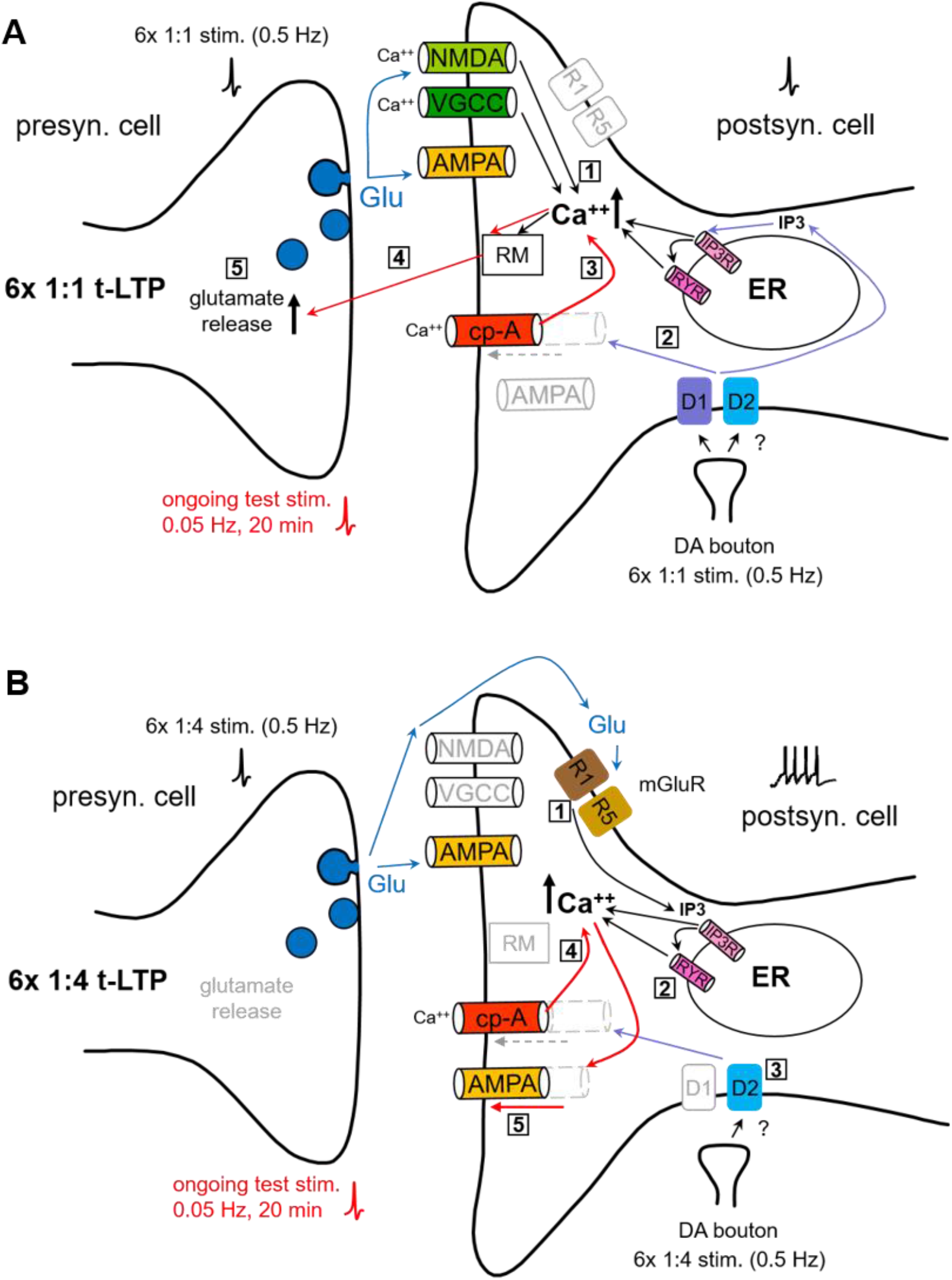
Suggested cellular signaling mechanisms involved in low repeat t-LTP at Schaffer collateral-CA1 synapses. Summary of induction and expression mechanisms involved in low repeat canonical (i.e. 6x 1:1) and burst (i.e. 6x 1:4) t-LTP protocols in CA1 pyramidal neurons. **A)** Prevailing presynaptic expression of 6x 1:1 t-LTP depends on postsynaptic induction via NMDAR and L-type VGCC mediated Ca^2+^ influx, supported by D1/D2 heterodimer induced Ca^2+^ release from the ER (1; compare text). Insertion of cp-AMPARs into the postsynaptic membrane might be regulated by D1/D2 signaling (2) and could account for the combined D1/D2 receptor dependence of 6x 1:1 t-LTP. Ongoing low frequency test stimulation after induction of t-LTP (shown in red) leads to sustained Ca^2+^ elevations through postsynaptic cp-AMPARs (3). The resulting prolonged postsynaptic Ca^2+^ elevation leads via a yet unidentified retrograde messenger (RM; BDNF, NO, endocannabinoids excluded) to increased presynaptic efficacy (4). An additional presynaptic contribution of D1/D2 signaling to enhanced presynaptic glutamatergic function is possible (not shown). **B)** The predominant postsynaptic expression of 6x 1:4 t-LTP does neither require postsynaptic NMDAR nor L-type VGCC activation for induction. It rather depends on Ca^2+^ release from postsynaptic internal stores mediated by mGluR_1,5_-dependent activation of IP3 receptors in the ER (1). This initial postsynaptic Ca^2+^ rise is amplified by Ca^2+^-dependent Ca^2+^ release via Ryanodine receptors (RyRs; 2). Moreover, the 6x 1:4 t-LTP depends (like 6x 1:1 t-LTP) on the activation of cp-AMPARs. Intact D2 receptor signaling is mandatory to observe 6x 1:4 t-LTP and might be involved in recruiting cp-AMPARs to the postsynaptic membrane (3) for sustained Ca^2+^ influx during ongoing low frequency test stimulation after t-LTP induction (4, shown in red). The resulting prolonged postsynaptic Ca^2+^ elevation initiated by mGluRs, RyRs, and cp-AMPARs leads to predominant postsynaptic expression of 6x 1:4 t-LTP by insertion of new GluA1/GluA2-containing AMPARs into the postsynaptic membrane (5.). The release of RM is supposedly inhibited by the different postsynaptic Ca^2+^ kinetics in response to the 6x 1:4 t-LTP compared to the 6x 1:1 protocol (compare Solinas et al., 2019).

## Discussion

Our study shows that t-LTP at hippocampal SC-CA1 synapses requires only six repeats of coincident presynaptic stimulation paired with either 1 or 4 postsynaptic spikes at low frequency (0.5 Hz). For the 1:4 burst protocol, even just three repeats are sufficient to elicit t-LTP. The 6x 1:1 t-LTP was induced by Ca^2+^ influx via postsynaptic NMDARs and L-type VGCCs, Ca^2+^ release from internal stores, and required combined D1/D2 receptor signaling. In contrast, the 6x 1:4 t-LTP was induced by postsynaptic Ca^2+^ release from internal stores mediated via mGluRs/IP_3_ signaling and ryanodine receptors, and was completely inhibited in the presence of D2 receptor antagonists. Both low repeat canonical and burst t-LTP occurred independent of BDNF release, but strongly depended on activation of GluA2-lacking cp-AMPARs. These data suggest that low repeat STDP paradigms can induce equally robust t-LTP as observed for high repeat t-LTP in the hippocampus. However, the pharmacological profile of low repeat t-LTP induction and expression revealed mechanistic differences between both induction protocols.

### Dependence of t-LTP on repeat number and frequency of the STDP stimulation

Both 6x t-LTP protocols used in our study yielded robust t-LTP with similar time courses as described previously for standard STDP paradigms that used either higher number of pairings or higher pairing frequency (compare e.g., Carlisle et al., 2008; Couey et al., 2007; Edelmann et al., 2015; Seol et al., 2007; Tigaret et al., 2016; Wittenberg and Wang, 2006; Yang and Dani, 2014). To date, only few studies focused on STDP protocols with low numbers of repeats for t-LTP induction (Cui et al., 2016; discussed in Edelmann et al., 2017; Froemke et al., 2006; Zhang et al., 2009). Since only such low repeat t-LTP protocols can be completed within a few seconds, these protocols are likely to represent a model for synaptic plasticity events triggering learning and memory processes that can also occur on a timescale of seconds. Thus, investigating the underlying signaling mechanisms might be relevant for learning induced synaptic changes *in vivo*. Similar to the results of Froemke and colleagues (Froemke et al., 2006) for layer 2/3 cortical neurons, we observed no significant difference in the magnitude of 1:1 t-LTP between the threshold repeat number (i.e., 6 repeats at 0.5 Hz) and higher repeat numbers at hippocampal Schaffer collateral (SC)–CA1 synapses (25 and 70 repeats; compare **Fig. 1A**). As for the canonical protocol, we also determined the threshold for successful t-LTP induction also for the burst protocol (compare **Fig. 1B**). The observed shift of the threshold repeat number to lower values (3 instead of 6 repeats for successful 1:4 t-LTP induction) for the burst protocol speaks in favor of facilitated postsynaptic induction by the spike train instead of single spikes used by the 1:1 protocol (compare Remy and Spruston, 2007). Together these data suggest that depending on the exact pattern (e.g., 1:1 vs. 1:4 paradigm) used for t-LTP induction distinct thresholds for the successful number of repeats can be observed.

Bittner and colleagues recently described in elegant *in vivo* recordings synaptic plasticity in mouse hippocampal place cells that can be triggered by pairing low numbers of postsynaptic action potentials with long-lasting dendritic depolarization, which works equally well with positive and negative pairing delays of roughly 1 s (Bittner et al., 2015; Bittner et al., 2017). While their work provides compelling evidence for the physiological relevance of low repeat spiking induced LTP for learning, this behavioral time scale synaptic plasticity follows a non-hebbian mechanism. In contrast, our low repeat t-LTP follows hebbian rules, since only simultaneous and nearly coincident pre- and postsynaptic pairing with short positive timing delays leads to associative potentiation (compare **Fig. 2**). Nevertheless, also such hebbian t-LTP protocols have been described previously to allow extension of STDP to behavioral time scales (compare e.g., Drew and Abbott, 2006; Gerstner et al., 2018; Shindou et al., 2019). In case of our low repeat t-LTP protocols, with the six repeat protocol comprising overall 10 s, and the three repeat protocol occurring within overall 4 s, this duration might bridge the time window between millisecond-dependent STDP and learned behavior on the time scale of several seconds.

In cultured hippocampal neurons, Zhang and colleagues (Zhang et al., 2009) showed that more than 10 repeats of their 1:1 STDP protocol were necessary to induce t-LTP. However, bath application of dopamine facilitated t-LTP induction and reduced the number of pairings that were required at a given frequency to successfully induce t-LTP (Zhang et al., 2009). Since primary cultures of dissociated hippocampal neurons develop synaptic connections in the absence of dopaminergic inputs, the role of endogenous DA can be investigated only if t-LTP is recorded in acutely isolated hippocampal slices as performed here. Interestingly, our data show that both low repeat t-LTP variants tested are blocked when signaling of endogenously released DA is inhibited (**Fig.7**). The release of endogenous DA in our slices (Edelmann and Lessmann, 2011, 2013) is therefore likely to account for the low number of repeats required for successful induction of t-LTP in our study. Whether this effect is due to acute release of DA from axon terminals elicited via the extracellular co-stimulation of dopaminergic afferents during t-LTP induction and test stimulation, or rather depends on ambient levels of DA in the slices, remains to be determined.

Regarding the magnitude of t-LTP induced by low repeat canonical and burst protocols, we found that both, 6x 1:1 and 6x 1:4 t-LTP, were equally successful to induce t-LTP at positive spike timings (**Fig. 2C**). Because it is reasonable to assume that 1:4 burst protocols induce longer lasting and stronger Ca^2+^ elevations than 1:1 pairings, it might be expected that the time course of synaptic potentiation could differ between the two protocols. However, both protocols induced t-LTP with comparable onsets and rise times of potentiation and also resulted in similar magnitudes of t-LTP after 1 h of recording (compare **Fig. 2C, D**). Thus, except for the lower threshold number of repeats to elicit t-LTP (see last paragraph), the burst protocol does not seem to be more effective in inducing t-LTP at SC-CA1 synapses than the canonical protocol. Future studies in reduced extracellular Ca^2+^, which was recently suggested to change mechanisms of synaptic plasticity (Inglebert et al., 2020), are required to clarify this issue for low repeat t-LTP.

We also compared different spike timings (with negative and positive delays), to compare the full capacity to induce bidirectional plasticity with low repeat protocols (**Fig. 2A, B**). For positive pairings with Δt: +20, +40, and +100 ms we observed a similar decline (compared to Δt: +10 ms) in t-LTP magnitude as described previously for higher numbers of repeats (compare Bi and Poo, 1998; Edelmann et al., 2015). When applying negative pairings (i.e. post before pre pairings) t-LTP was absent, but we did not observe robust t-LTD for either of the two protocols. While these results stress that successful induction of t-LTP is critically dependent on the sequence of presynaptic and postsynaptic spiking and on the pairing interval, future studies should address under which conditions low repeat t-LTD can be induced by anti-causal synaptic activation.

### Mechanisms of expression of low repeat t-LTP

Despite the similarities described above, both low repeat protocols seem to recruit different expression mechanisms. For potentiation induced with the 6x 1:1 protocol, presynaptic mechanism seem to prevail (see below), whereas the 6x 1:4 protocol relies predominantly on postsynaptic insertion of AMPA receptors (**Fig. 3**). Commonly, LTP at SC-CA1 synapses induced by high-frequency stimulation is also thought to be expressed by a postsynaptic increase in AMPA receptor mediated currents (Granger and Nicoll, 2014; Nicoll, 2003). For STDP, however, different mechanisms of expression have been described that varied between brain regions and depending on experimental conditions (see e.g., Costa et al., 2017). Even at a given type of synapse (i.e. hippocampal SC-CA1) t-LTP can be expressed either pre- or postsynaptically (Edelmann et al., 2015). At this synapse, the expression mechanism of LTP seemed to be encoded by the pairing pattern used for STDP. While t-LTP induced by 70x 1:1 stimulation was expressed via increased presynaptic glutamate release, a 35 × 1:4 t-LTP was expressed via insertion of additional AMPARs by a GluA1-dependent mechanism (Edelmann et al., 2015). However, in this previous study, we used different numbers of repeats for the two t-LTP protocols (i.e. 20-35 × 1:4 and 70-100 × 1:1) to keep postsynaptic activity at an equivalent level. Those previous results did not allow to distinguish whether repeat number or stimulation pattern determined the site of t-LTP expression. With help of our current experiments using fixed numbers of repeats for both protocols, we could now determine that the pattern of postsynaptic spiking and not the repeat number influences the expression locus for t-LTP (compare **Fig. 3**).

For the 6x 1:1 t-LTP, the absence of an increase in AMPAR mediated currents and the independence from insertion of new AMPA receptors (**Fig. 3D, E**) speak against a strong contribution of postsynaptic expression, whereas the observed significant decrease in paired pulse ratio (PPR) and the increased mEPSC frequencies after successful induction of t-LTP (**Fig. 3A, B**), are consistent with a prevailing presynaptic enhancement of glutamate release probability. Regarding the retrograde messenger required for 6x 1:1 t-LTP, our data indicate that neither BDNF (**Fig. 8**) nor NO or endocannabinoids (Fig. S1) are involved in the presynaptic expression. However, further investigating the underlying presynaptic mechanisms of 6x 1:1 t-LTP was beyond the scope of the current study.

The expression mechanism for the six repeat version of our burst t-LTP protocol (6x 1:4) seems to follow the suggested mechanisms for conventional SC-CA1 LTP, with postsynaptic expression via insertion of new AMPARs leading to an increased AMPAR/NMDAR mediated current ratio (**Fig. 3D**). Such a postsynaptic mechanism is also favored by the increased amplitudes of mEPSCs following successful induction of 6x 1:4 t-LTP (**Fig. 3C**). - The absence of a significant change in paired pulse facilitation and the lack of an increase in mEPSC frequencies after successful induction of 6x 1:4 t-LTP speak against a strong presynaptic contribution for low repeat burst t-LTP (**Fig. 3A, C**). Most importantly, our experiments with Pep1-TGL clearly demonstrate the importance of GluA1 containing AMPARs for the expression of 6x 1:4 t-LTP (**Fig. 3E**) while 6x 1:1 t-LTP is completely independent from this manipulation.

### Dependence of low repeat t-LTP induction on different sources for postsynaptic Ca^2+^ elevation

Investigating the mechanisms of t-LTP induced with low repeat STDP protocols has just started. Accordingly, the contribution of different sources of Ca^2+^ to its induction was until now largely unknown. Unexpectedly, our experiments revealed distinctly different routes for postsynaptic Ca^2+^ elevation for the low repeat 1:1 and 1:4 protocols to induce t-LTP. The results for the 6x 1:1 t-LTP are in accordance with previous studies showing that t-LTP as well as classical high frequency stimulation induced LTP at CA1 glutamatergic synapses rely on Ca^2+^ influx via postsynaptic NMDA receptors (Malenka and Bear, 2004). For STDP, NMDARs are thought to serve as coincidence detectors of timed pre- and postsynaptic activation (e.g., Bi and Poo, 1998; Debanne et al., 1998; Edelmann et al., 2015; Feldman, 2000). Depending on the level of postsynaptic Ca^2+^ that is reached during induction, separate signaling cascades leading to either LTP or LTD are activated (Artola and Singer, 1993; cited in Caporale and Dan, 2008; Lisman, 1989). For t-LTD, alternative mechanisms for coincidence detection have been described (Bender et al., 2006; Fino and Venance, 2010). Instead of NMDAR-mediated Ca^2+^ influx, these studies reported that either mGluRs, L-type VGCCs or IP3 gated internal Ca^2+^ stores can trigger the induction of LTD. As for LTD, also for LTP, additional coincidence detectors and Ca^2+^ sources might be involved in its induction (Dudman et al., 2007; VGCC: Magee and Johnston, 1997; Nanou et al., 2016; IP3-sensitive stores: Takechi et al., 1998; Wang et al., 2016; Wiera et al., 2017). In accordance with these previous studies, we found that 6x 1:1 t-LTP can in addition to NMDARs also be induced by Ca^2+^ entry through L-type VGCCs (**Fig. 4C, E**), and is supported by IP3- and RyR-dependent Ca^2+^ release from internal stores (**Fig. 6B, C**).

In contrast to these conventional Ca^2+^ sources for the canonical low repeat t-LTP, the situation is much different for 6x 1:4 burst t-LTP. Although a requirement for postsynaptic Ca^2+^ elevation is clearly evident from the BAPTA experiments (**Fig. 4B**), Ca^2+^ entry via NMDARs or L-type VGCCs was not involved (compare **Fig. 4D, F**). Rather, our results demonstrated that the initial postsynaptic Ca^2+^ rise involved group I mGluRs (i.e. mGluR1 and mGluR5; Kaar and Rae, 2015), subsequent activation of IP_3_Rs and RyRs (compare **Fig. 5**), eventually activating (like the 6x 1:1 protocol) GluA2-lacking Ca^2+^-permeable AMPARs in the postsynaptic membrane (**Fig. 10**). While activation of mGluRs seems to contribute to the initial postsynaptic Ca^2+^ rise in 6x 1:4 t-LTP, subsequent Ca^2+^ induced Ca^2+^ release via RyRs amplifies and prolongs this Ca^2+^ signal (compare **Fig. 5D**). The initial rise in postsynaptic Ca^2+^ levels might be co-induced by Ca^2+^ influx through GluA2 subunit deficient Ca^2+^-permeable AMPA receptors (cp-AMPARs) into the postsynaptic cell (Suzuki et al., 2001). This overall interpretation is concluded from our experiments performed in the presence of the mGluR antagonists, IP_3_R inhibitors and the antagonists of Ca^2+^ permeable AMPARs, NASPM and IEM, which completely inhibited 6x 1:4 t-LTP (group I mGluR: **Fig. 5A, B**, cp-AMPAR: **Fig. 9B, D**, for discussion of cp-AMPAR, see below).

Group I metabotropic GluR have indeed been described previously to contribute to certain types of hippocampal LTP (Wang et al., 2016), while our present results show for the first time their involvement in STDP. Altogether it seems plausible that 6x 1:4 stimulation first activates mGluR_1,5_ receptors, which subsequently trigger IP_3_ mediated calcium release from internal stores (Jong et al., 2014, compare Fig 5D) compare **Fig. 5C**). The resulting calcium rise and additional Ca^2+^ influx via cp-AMPARs might then be strengthened by additional IP_3_ and RyR mediated calcium induced Ca^2+^ release to successfully boost low repeat induced burst t-LTP (compare **Fig. 10**).

### Regulation of low repeat t-LTP by dopamine receptor signaling

Since high repeat STDP is regulated by dopamine (e.g., Cassenaer and Laurent, 2012; Cui et al., 2015; Edelmann et al., 2015; Edelmann et al., 2017; Edelmann and Lessmann, 2011, 2018; Pawlak and Kerr, 2008; Seol et al., 2007; Yang and Dani, 2014; Zhang et al., 2009), we also investigated DAergic modulation of our two low repeat STDP variants (compare **Fig. 7**). The 6x 1:4 t-LTP was dependent entirely on intact D2 receptor signaling This result can be easily reconciled with pure D2 receptor-dependent signaling being responsible for induction of 6x 1:4 t-LTP (compare **Fig. 7B**). Little is known about D2R mediated function in t-LTP and classical LTP. It was shown, however, that D2 receptors can limit feedforward inhibition in the prefrontal cortex and allow thereby more effective t-LTP (Xu and Yao, 2010). Importantly, D2-like receptors are expressed in the hippocampus in pre- and postsynaptic neurons and have been described to regulate synaptic plasticity (Beaulieu and Gainetdinov, 2011; Dubovyk and Manahan-Vaughan, 2019; Sokoloff et al., 2006). Moreover, D2 receptors contribute to hippocampus-dependent cognitive functions (Nyberg et al., 2016). Together, these previous results on D2 receptor functions are in line with the role in 6x 1:4 t-LTP. In contrast to the 6 repeat burst protocol, the 6x 1:1 t-LTP remained functional when either D1-like or D2-like dopamine receptor signaling was intact. Moreover, while the D1 receptor inhibitor SCH23390 alone did not show any signs of 6x 1:1 t-LTP inhibition, it was nevertheless able to impair the slightly reduced t-LTP in the presence of Sulpiride down to control levels, when both antagonists were co-applied (**Fig. 7A**). The interpretation of this pharmacological profile of 6x 1:1 t-LTP needs to take into consideration that D1-like and D2-like receptors do not signal exclusively via altering cAMP levels (cAMP increase via D1-like receptors - or decreased via D2-like receptors; Tritsch and Sabatini, 2012). Rather, D5 receptors and heterodimeric D1/D2 receptors can also activate PLC pathways stimulating in turn IP_3_/Ca^2+^, DAG/PKC signaling, or MAPK signaling downstream of D1 receptors. Also direct modulation of NMDARs and VGCCs in response to D2R activation is possible (Beaulieu and Gainetdinov, 2011). Whether the combined D1-like/D2-like receptor dependence of 6x 1:1 t-LTP reflects indeed activation of D1/D2 heteromers needs to be addressed by future experiments.

To interpret the combined regulation of the 6x 1:1 t-LTP by D1- and D2-like receptors it also needs to be taken into consideration that D2-like receptors are generally believed to display a higher affinity for DA compared to D1-like receptors (Beaulieu and Gainetdinov, 2011). This co-regulation could assure that slowly rising ambient DA levels created by tonic firing of DAergic neurons are equally effective in regulating 6x 1:1 t-LTP as much faster rising DA concentrations during phasic firing.

Such a change in DA release was indeed shown in recordings of midbrain neurons, where activity of DAergic neurons switches from tonic to phasic burst activity resulting in locally distinct levels of secreted DA in the target regions (Rosen et al., 2015). Local DA concentration differences can then result in different DA-dependent effects, with high affinity D2-like receptors being activated by low and slowly rising extracellular DA levels, while low affinity D1 receptors are only activated by local DA peaks. In our STDP experiments, where DAergic input fibers are most likely co-activated during SC stimulation, we observed similar activity-dependent recruitment of different DA receptors. While D1 receptor-dependent effects were activated by 70-100x 1:1 stimulation (Edelmann and Lessmann, 2011 and compare **Fig 7C**), D1/D2 receptors or pure D2 receptor mediated processes were already activated by six presynaptic co-stimulations of DAergic fibers (compare **Fig. 7A, B**). Taking into account that D2-like receptors (i.e. D2, D3 and D4 receptors) are classically thought to inhibit LTP by decreasing cAMP/PKA signaling (Otmakhov and Lisman, 2002; Otmakhova et al., 2000), D2-like receptor driven processes promoting t-LTP might indeed be activated by G_βγ_ signaling independent of cAMP pathways. G_βγ_ signaling also blocks L-type and N-type VGCCs (Tritsch and Sabatini, 2012) and D2 receptor signaling can yield Ca^2+^ release from internal stores - two mechanisms that might account for the uncommon type of calcium source required for the induction of our 6x 1:4 t-LTP (compare **Figs. 4, 5, 10**).

### Independence of low repeat t-LTP from BDNF/TrkB signaling

Brain-derived neurotrophic factor (BDNF) is well known for its important role in mediating long-lasting changes of synaptic plasticity (reviewed in e.g., Edelmann et al., 2014; Gottmann et al., 2009; Lessmann et al., 2003; Park and Poo, 2013). Moreover, BDNF is also involved in regulating STDP (Edelmann et al., 2015; Lu et al., 2014; Sivakumaran et al., 2009). For hippocampal SC-CA1 synapses it was shown that BDNF is secreted from postsynaptic CA1 neurons in response to 20-35 repeats of a 1:4 STDP protocol mediating postsynaptically expressed t-LTP via postsynaptic TrkB receptor activetion (Edelmann et al., 2015). Interestingly, the results of the present study revealed, that neither of the two low repeat t-LTP variants depends on BDNF induced TrkB signaling (compare **Fig. 8**). This finding was not unexpected since release of endogenous BDNF has been reported previously to require more prolonged barrages of AP firing than just 6 repeats of short AP (burst) firing at 0.5 Hz (compare Balkowiec and Katz, 2002; Edelmann et al., 2015; Lu et al., 2014). This BDNF independency was observed in situations with either chronic (e.g., heterozygous BDNF ko animals) or acute depletion of BDNF (BDNF scavenger; see e.g., Edelmann et al., 2015; Meis et al., 2012; Schildt et al., 2013).

### Function of GluA2-lacking Ca^2+^ permeable AMPA receptors in low repeat t-LTP

Interestingly, both variants of low repeat t-LTP were strictly dependent on activation of GluA2-lacking calcium-permeable (cp-) AMPA receptors (**Fig. 9**). In the respective experiments, NASPM or IEM were present in the ACSF from the start of the recording to assure complete inhibition of cp-AMPARs during t-LTP induction. In CA1 neurons, cp-AMPARs were described to be absent from postsynaptic membranes during basal synaptic stimulation. Rather, they were reported to transiently insert into the postsynaptic membrane after tetanic LTP stimulation to allow sustained Ca^2+^ influx into the postsynaptic neuron after LTP induction, thereby facilitating expression of late LTP (reviewed in Park et al., 2018). A role of cp-AMPARs in STDP has thus far not been reported and these results represent a crucial new finding that emerges from our study. Additional experiments will be required to determine the time course of activity-dependent cp-AMPAR incorporation during induction of low repeat t-LTP into the postsynaptic membrane. Furthermore, it needs to be determined how cp-AMPAR mediated Ca^2+^ influx is orchestrated with mGluR- and RyR-dependent Ca^2+^ elevation for induction of low repeat 6x 1:4 t-LTP. Likewise, the co-operation of cp-AMPARs with NMDAR- and VGCC-dependent Ca^2+^ elevations for inducing 6x 1:1 t-LTP needs to be investigated.

In addition to allowing sufficient Ca^2+^ elevation in t-LTP, cp-AMPARs might be involved in DA-dependent priming of synapses for delayed/retroactive reinforcement of LTP or silent eligibility traces (e.g., Brzosko et al., 2015; Gerstner et al., 2018; He et al., 2015; Shindou et al., 2019). By those eligibility traces or delayed reinforcements, the different time scales between milliseconds and seconds can be bridged, thereby allowing to connect hebbian synaptic plasticity to behavioral responses and learning. Such mechanisms might also be involved in the signaling mechanisms employed by our low repeat t-LTP protocols (6x 1:1 and 6x 1:4), since both variants of t-LTP show a clear dependence on DA signaling and on cp-AMPARs (compare **Figs. 7** and **9**).

In summary, we used two different low repeat STDP protocols at SC-CA1 synapses to record synaptic plasticity at the single cell level in postsynaptic CA1 neurons (i.e. t-LTP). We found that, dependent on stimulation pattern and repeat number, distinct signaling and expression mechanisms are activated by the canonical and the burst low repeat paradigm. From our experiments, we can conclude that even with the same experimental setup, age and species, multiple types of synaptic plasticity mechanisms can coexist at a given type of synapse. This plethora of coexisting plasticity mechanisms for strengthening synaptic transmission seems to be ideally suited to empower the hippocampus to fulfill its multiplexed functions in memory storage.

## Acknowledgments

The project was funded by the federal state of Saxony-Anhalt and the “European Regional Developmental Fund (ERDF 2014-2020), Project: Center for Behavioral Brain Sciences (CBBS) FKS: ZS/2016/04/78113; by the federal state Saxony-Anhalt and the European Structural and Investment Funds (ESF, 2014-2020), project number ZS/2016/08/80645 (ABINEP) and by the DFG (ED 280/1-1 and SFB779/B06). The authors thank Regina Ziegler for excellent technical assistance.

CRediT

Conceptualization: VL, EE; Methodology: VL, EE and EC-P; Investigation: EC-P, BK, GQ, SB with help by EE; Writing: EE, VL, with help by EC-P; Funding Acquisition: VL, EE; Resources: VL; Supervision: VL, EE.

The authors declare no competing interests.

---

**Figure S1:**
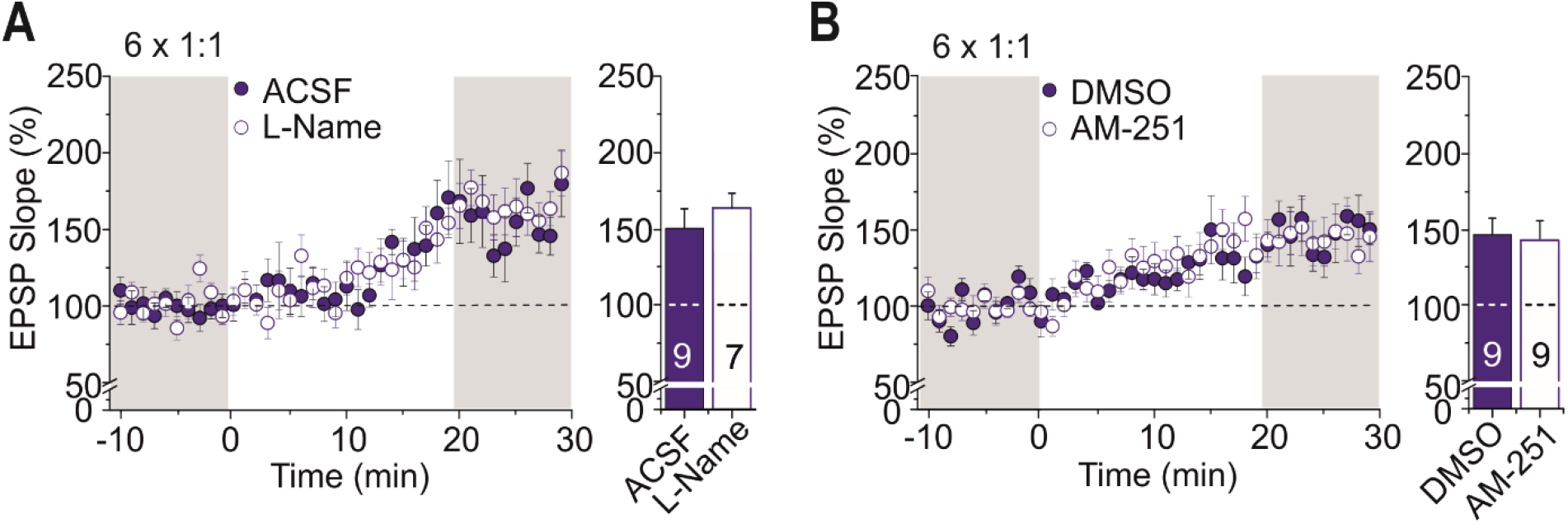
6x 1:1 t-LTP does not depend on nitric oxide (NO) or endocannabinoid signaling via CB1 receptors. SC-CA1 synapses were recorded as in **figure 4**, and 6x 1:1 t-LTP stimulation was performed at 0 min. **A)** NOS signaling was inhibited by bath application of L-NAME (100 µM) but did not reveal inhibition of 6x 1:1 t-LTP. **B)** Inhibition of CB1 receptors with AM-251 (3 µM) did not inhibit 6x 1:1 t-LTP.

**Figure S2:**
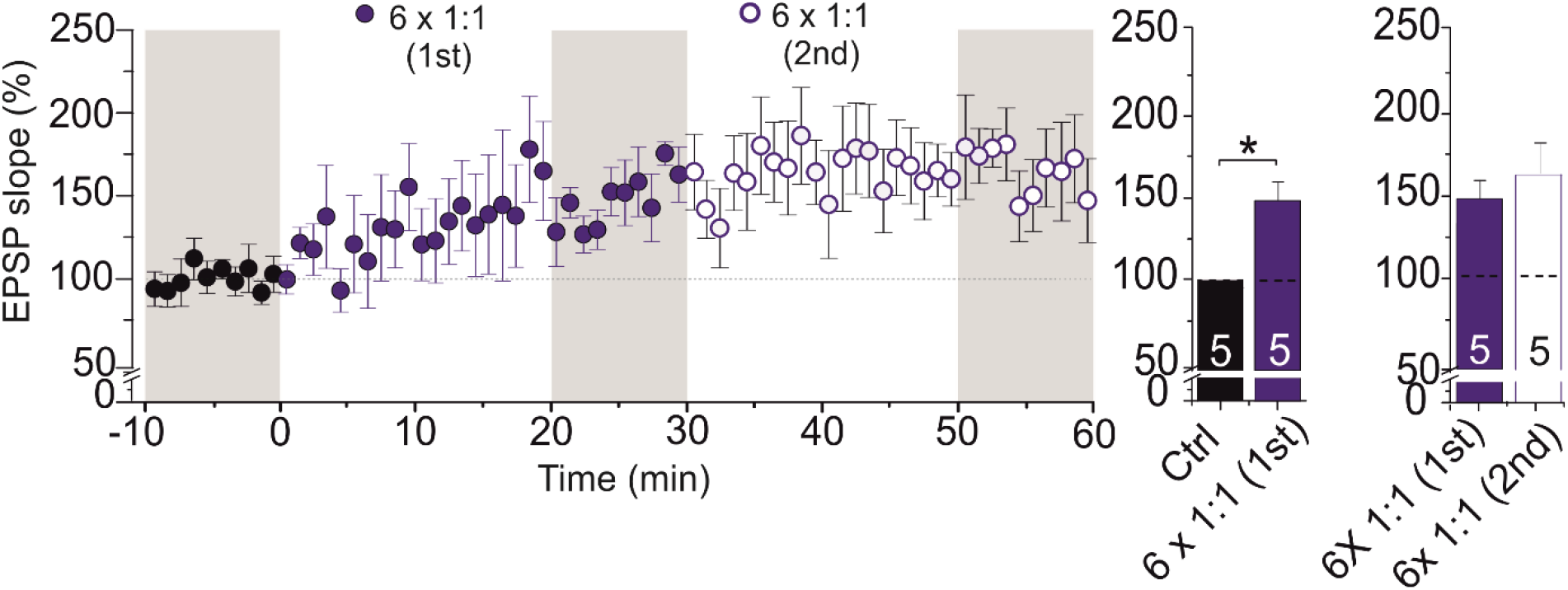
Two subsequent stimulations with the 6x 1:1 t-LTP protocol do not yield additional potentiation. SC-CA1 synapses were recorded as in **figure 9**, and 6x 1:1 t-LTP stimulation was performed at 0 and 30 min in the same cells (n=5 / N= 3). The second induction protocol did not significantly increase the magnitude of t-LTP that was reached after the first t-LTP induction. Note that subsequent stimulations with the 6x 1:1 protocol followed by the 6x 1:4 protocol in the same cells yielded additional and independent potentiation (compare **Fig. 9E**). Average time course of potentiation and mean (± SEM) magnitude of t-LTP are shown for the respective experiments.

**Figure S3:**
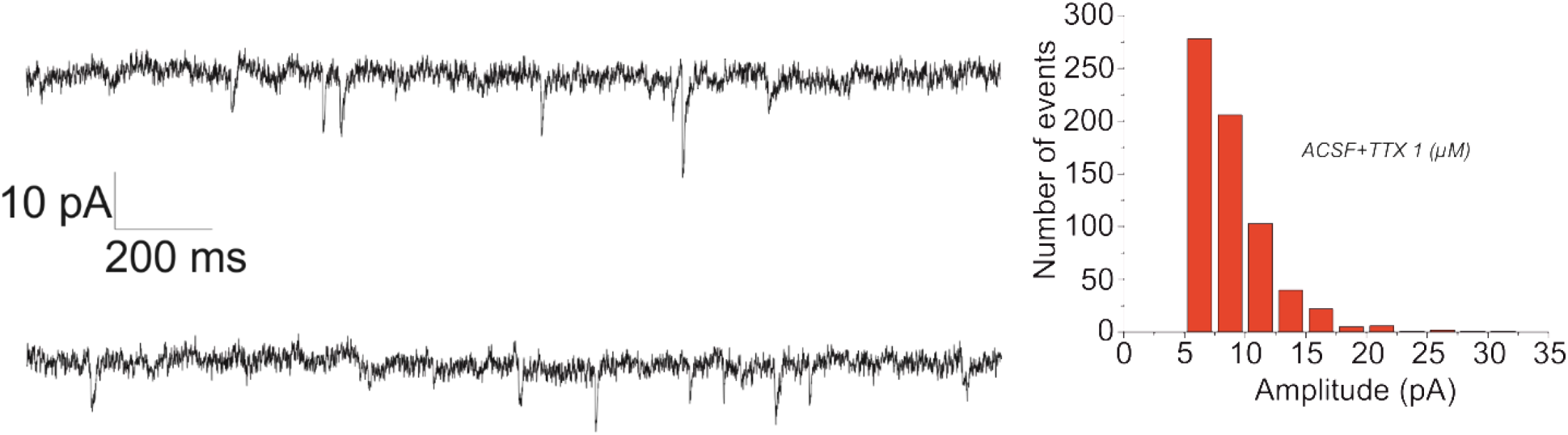
Miniature (m)EPSC recorded in the presence of 1 µM TTX and 50 µM picrotoxin in ACSF recording solution. **Left:** Original traces of mEPSCs recorded in voltage clamp at -70mV holding potential. **Right:** Number of mEPSC events during 5 minutes of recording from 4 different CA1 pyramidal cells under the same conditions as used in t-LTP experiments. Results show that 99% of mEPSCs are <20 pA, therefore justifying our cut-off frequency.

